# A primordium-level ice barrier in *Vaccinium corymbosum* floral buds is lost as their xylem trace matures

**DOI:** 10.64898/2026.10.29.762774

**Authors:** Camilo Villouta, Ava K. Russell, Angelique Acevedo

**Author notes:** Correspondence: Camilo Villouta.

## Abstract

Flower primordia of *Vaccinium corymbosum* L. (highbush blueberry) exclude ice formation in winter through an unexplained biological barrier despite lignified xylem connecting them to the stem. We asked where this barrier lies and whether the maturation of that connection, rather than its presence, tracks its seasonal loss. On buds collected in fall, winter and spring we combined structure-resolved freezing tests, water content, thermal analysis, cryomicroscopy and xylem anatomy. Whole-bud LT_50_ was -20 °C in fall, -31 °C in winter and -16 °C in spring; water content rose only in spring. Buds froze extraorgan, without low-temperature exotherms to -46 °C. Ice nucleated in the stem, reached the bud within minutes and accumulated in the scales. The bud axis froze in every season, whereas the primordia stayed ice-free in fall and winter and froze in spring, placing the barrier at each primordium, not the bud base. A continuous xylem trace reached every primordium in every season; its elements were mostly annular and helical until spring, when scalariform, reticulate and pitted elements predominated. Junction conduits stayed narrow (2 to 3 µm). The primordium-level barrier was lost as this trace matured; we propose that its immature state is part of the long-unexplained barrier.

## Introduction

Floral buds are among the most freeze-sensitive organs of temperate woody perennials, and the depth and timing of their freezing tolerance decide reproductive success and, in fruit crops, yield (Rodrigo 2000). Hardiness is gained during fall acclimation, peaks in midwinter and is lost during spring deacclimation (Kalberer et al. 2006). Risk therefore concentrates at the two transitions, and warmer, more variable late winters shift it rather than remove it (Arora & Taulavuori 2016; Zohner et al. 2020; Kirchhof et al. 2025; Charrier et al. 2026). *Vaccinium corymbosum* L. (highbush blueberry) is a good system for asking what sets this pattern, because the pattern itself is well described. Cold hardiness is measured in controlled freezing tests as the temperature that kills a given fraction of the flower primordia (Bittenbender & Howell 1974, 1976; Rowland et al. 2005), and the midwinter LT_50_ of hardy northern highbush cultivars reaches about -25 to -30 °C in the field (Ehlenfeldt et al. 2007; Hanson et al. 2007; Rowland et al. 2008). The kinetics of deacclimation, models of hardiness and the molecular acclimation program are also known (Muthalif & Rowland 1994; Arora et al. 2004; Arora & Rowland 2011; Kogan et al. 2023). What is not known is its anatomical basis: where ice forms in the bud and what keeps it out of the primordia.

Overwintering buds survive freezing in two ways, and both depend on keeping ice out of the primordia (Burke et al. 1976; Sakai & Larcher 1987; Wisniewski et al. 2014). In deep supercooling, the primordial water stays liquid far below 0 °C until it freezes; that freezing is lethal, and the heat it releases can be recorded by differential thermal analysis (DTA) and be described as a low-temperature exotherm (LTE), as in *Prunus* and *Rhododendron* (George et al. 1974; Ishikawa & Sakai 1981; Quamme et al. 1995). In extraorgan freezing, the primordia stay ice-free by losing water to ice masses in the surrounding bud scales and axis, as in *Cornus*, larch and many vegetative buds (Ishikawa & Sakai 1985; Endoh et al. 2014; Neuner et al. 2019a). *V. corymbosum* belongs to the second group, on two grounds. LTEs have been recorded in its floral buds (Biermann et al. 1979) but do not predict injury (Flinn & Ashworth 1994a), so injury is not marked by an exotherm and hardiness is measured by controlled freezing tests rather than by DTA. Direct observation agrees: in buds fixed while frozen, ice formed in the scales and bracts at every stage, whereas the rachises, pedicels and florets held no ice while the buds were hardy and froze only in nonacclimated and deacclimated buds (Flinn & Ashworth 1994b). Cranberry (*V. macrocarpon*) behaves the same way, with no LTE, freeze dehydration seen by magnetic resonance imaging, and a temporary ice barrier at the bud base in fall and winter only (Villouta et al. 2020, 2021, 2022). In both species the ice comes from the stem: ice nucleators in *Vaccinium* stem bark initiate freezing at -1 to -4 °C (Kishimoto et al. 2014), and ice propagates from the stem into attached leaves and flowers (Wisniewski et al. 1997; Workmaster et al. 1999). The question is what stops it between stem and primordia.

Two anatomical models explain how ice is kept out of the primordia. In the first, the barrier is the absence of a conduit, like xylem. In *Prunus* species whose flower buds deep supercool, the dormant primordia have procambial strands but no differentiated xylem, whereas species whose buds do not supercool have continuous vessels (Ashworth 1984; Kader & Proebsting 1992). In spring, vessels differentiate acropetally into the primordia as the capacity to supercool is lost, as in *Forsythia*, apricot and sweet cherry (Ashworth et al. 1989, 1992; Andreini et al. 2012; Houghton et al. 2024). In the second model the xylem connection is intact and other features stop the ice: a constriction of tracheids at the pedicel base lets *Calluna vulgaris* reproductive shoots supercool below -20 °C (Kuprian et al. 2014, 2016; Bertel et al. 2021), permeability barriers block ice in the bud axis of *Vitis* (Jones et al. 2000), and bud architecture and wall chemistry isolate *Picea* primordia (Kuprian et al. 2017). *V. corymbosum* fits neither model. Flinn and Ashworth (1994b) tested the first directly: cleared buds stained for lignified xylem showed continuous strands from rachis through pedicels into the lower corolla in nonacclimated and acclimated buds alike, stagnant from December to February and proliferating throughout the florets by mid-April. Because ice stayed out of florets connected to the stem by xylem, they concluded that the barrier is something other than a missing conduit, and left its nature open. A lignin stain, however, does not report differentiation state. Xylem differentiates in stages: in a growing organ, protoxylem elements with annular or helical thickenings form first and metaxylem elements with scalariform, reticulate or pitted walls later, and both are lignified (Evert 2006). A trace could therefore be continuous and lignified yet not fully differentiated, and whether its differentiation state changes over the dormancy cycle has not been examined in *V. corymbosum*.

The junction matters too, because conduit anatomy bears on freezing in three ways. First, ice travels fastest along vascular tissue (Pearce & Fuller 2001; Hacker & Neuner 2007; Neuner et al. 2010), and its passage between conduits depends on the openings it crosses: water in small pores freezes only at depressed temperatures (Ashworth & Abeles 1984), and pit membranes and wall modifications restrict ice propagation (Wisniewski et al. 1991; Takahashi et al. 2021). Second, conduit diameter sets vulnerability to freeze-thaw embolism (Sperry & Sullivan 1992; Davis et al. 1999; Pittermann & Sperry 2003, 2006; Mayr & Sperry 2010). Third, wall thickness relative to lumen span, (*t*/*b*)², sets resistance to collapse under negative pressure (Hacke et al. 2001). In cranberry, ice stops in fall and winter at a bud base crossed by tracheids that kept their diameter from September to May but increased in number as xylogenesis resumed in spring, and were proposed to be non-functional until then (Villouta et al. 2022). To our knowledge no comparable seasonal measurements exist for *V. corymbosum*.

What is missing for *V. corymbosum* is a seasonal account in which hardiness, the location and timing of ice, and the anatomy of trace and junction are measured on the same material. Our objectives were to determine, in floral buds collected in fall, winter and spring, (i) bud water content and structure-resolved hardiness, by controlled freezing tests; (ii) where and when ice forms, by infrared differential thermal analysis (IDTA), DTA and reflected-polarised-light cryomicroscopy; (iii) the presence and differentiation state of the xylem trace to the primordia, by light microscopy with wall-pattern scoring; and (iv) the conduit anatomy of the bud base and subtending stem. We hypothesised that the state of the vascular connection between stem and primordia, the differentiation of the trace and the conduit anatomy of the junction, helps set the seasonal envelope of floral-bud freezing tolerance, with bulk dehydration a consequence, not a cause. If so, mean bud water content should not track LT_50_; freezing should start in the stem and reach the bud after a delay, without an LTE; the trace should mature from protoxylem-type to metaxylem-type elements only in spring; and the bud base, but not the stem, should shift in winter towards more numerous, narrower and thicker-walled conduits. We also expected our material to reproduce the pattern of Flinn and Ashworth (1994b): hardiness deepening to midwinter and lost by spring, ice in the scales with ice-free primordia in fall and winter, and a xylem trace reaching the primordia in every season.

The answers matter for the crop and beyond it. Cultivars differ in susceptibility to spring frost (Hancock et al. 1987), and cultivar-specific models that predict lethal temperatures can guide freeze protection (Kogan et al. 2023). A lethal temperature says how much cold the primordia survive, not why; a structural trait that tracks the loss of ice exclusion could give these models a mechanism and breeders a marker. Beyond the crop, comparison with *V. macrocarpon* asks whether the barrier lies at the bud base in both congeners. More generally, *V. corymbosum*, which fits neither model, can show whether the spring reconnection of bud and stem may occur by maturation of a trace that was never absent.

## Materials and methods

### Plant material and sampling

Floral buds of *V. corymbosum* were collected from five established plants at the Arnold Arboretum of Harvard University (Boston, MA, USA; 42.28° N, 71.13° W), Arboretum accessions 22898*A, 20745*A, 20747*A, 20748*A and 250-34*A, on three dates, used for every assay, spanning the dormancy cycle: 25 October 2021 (fall, acclimating), 20 January 2022 (winter, maximum hardiness) and 23 March 2022 (spring, deacclimating). Bud-bearing stem segments were cut from the five plants on each date, sealed in bags with moist paper and carried to the laboratory on ice, stored at 4 °C and processed within 24 h. ‘Floral bud’ denotes the whole organ (scales, bud axis, rachis and flower primordia) and ‘flower primordia’ the individual florets. The same plants were sampled on every date, and each assay drew its buds from all five, or from as many as its sample size allowed; plant identity was not retained in the analyses.

### Bud water content

Twenty buds per collection were excised at the bud-stem union, weighed fresh and dried at 65 °C for 72 h to constant mass. Water content was expressed as a percentage of fresh mass (Figure 1A).

**Figure 1.**
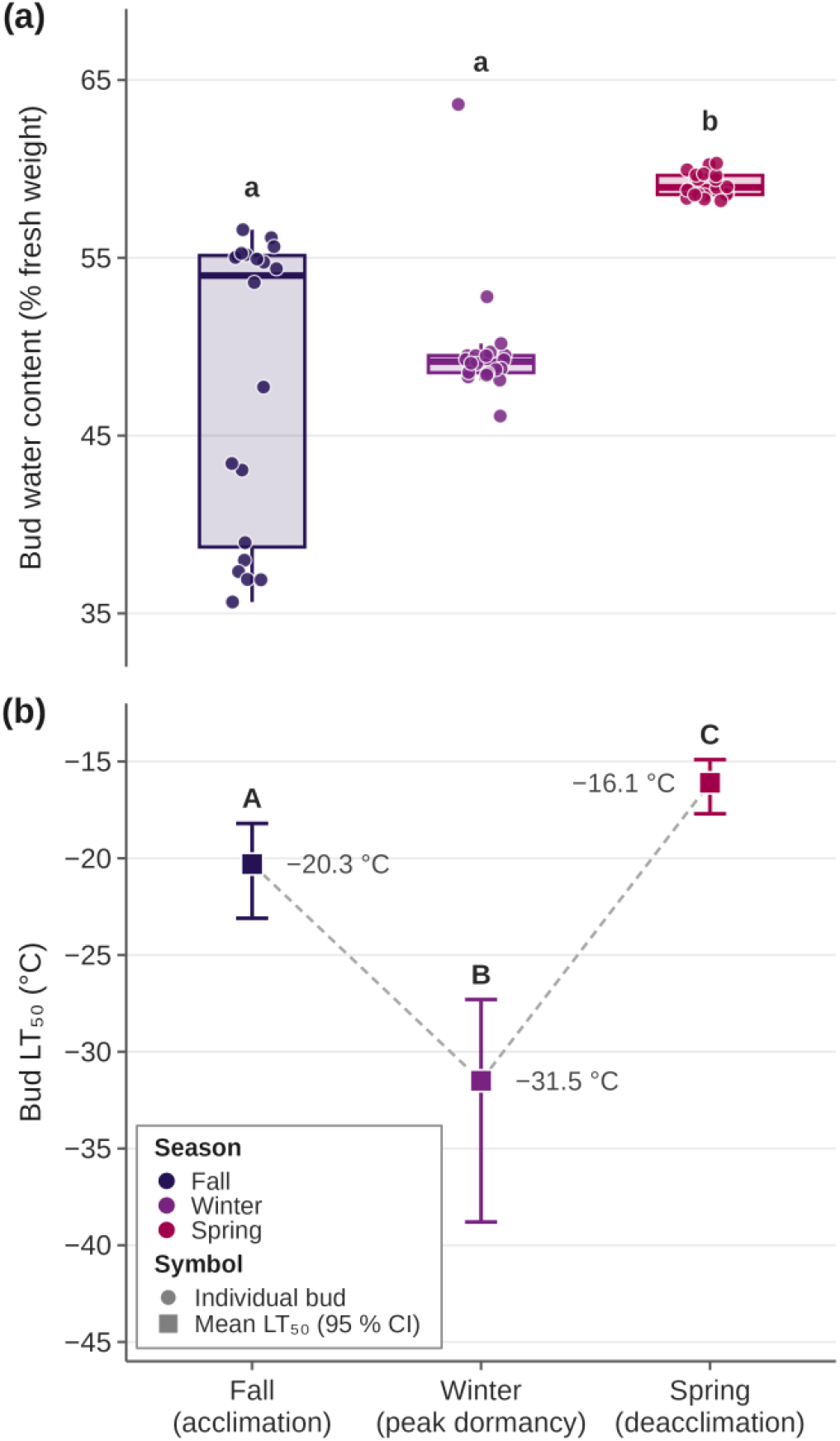
Bud water content and whole-bud freezing tolerance of *Vaccinium corymbosum* floral buds across the dormancy cycle. (A) Water content (% of fresh mass) of individual buds (points; *n* = 20 per season); boxes show median, interquartile range and 1.5 x IQR whiskers; different lowercase letters denote seasons that differ (Kruskal-Wallis, pairwise Wilcoxon with Holm correction, *P* < 0.05). (B) Whole-bud LT_50_ from the controlled freezing tests of Figure 2 with 95 % Monte Carlo confidence intervals; different uppercase letters denote non-overlapping intervals; the winter interval is extrapolated (non-saturating curve). Seasons: fall 2021, winter 2022, spring 2022; season colours are the same in all figures.

### Controlled freezing tests and lethal temperatures

Controlled freezing tests followed Villouta et al. (2020). Twigs about 8 cm long with at least one floral bud were rinsed, blotted dry, wrapped at the base in moist paper towel to prevent artificial supercooling and placed in 50 ml centrifuge tubes, three tubes per target temperature holding three, three and four twigs (ten buds per temperature). Tubes were equilibrated at 4 °C for 3 h in a programmable freezing chamber (Tenney Junior, Thermal Product Solutions, New Columbia, PA, USA) cooled to -1 °C at 1 °C h⁻¹, held for 1 h to promote ice nucleation, then cooled at 1 °C h⁻¹ to -6 °C, 2 °C h⁻¹ to -12 °C and 4 °C h⁻¹ to -50 °C. Tubes were removed at -4, -8, -12, -16, -20, -24, -32, -40 and -50 °C, with an unfrozen control at 4 °C, kept on ice in the dark for 12 h, at 4 °C for 3 d and at room temperature in low light for 24 h to let injured tissue brown. Buds were cut longitudinally and examined under a stereomicroscope (Discovery V12, 1.5x PlanApo objective, Axiocam 512 color camera; Zeiss, Jena, Germany). Injury was scored from 0 (no browning) to 3 (complete browning and collapse) for the subtending stem, the bud scales, the bud axis and the flower primordia with the shoot apical meristem, on the scale used for cranberry buds (Villouta et al. 2021); the whole-bud score was the mean of the four structure scores.

Mean injury per temperature was fitted, for each season and structure and for the whole-bud mean, to a three-parameter Gompertz function, injury = *A* exp[-exp(*b* - *c*|*T*|)], with *A* the upper asymptote, *b* the location, *c* the rate and |*T*| the absolute value of the treatment temperature (Lim et al. 1998), by nonlinear least squares (nlsLM, minpack.lm; Elzhov et al. 2023) and bounds *A* in [0.5, 3.5], *b* in [-5, 10], *c* in [0.001, 5]. Lethal temperatures were obtained analytically as LTₚ = -[*b* - ln(-ln *p*)]/*c* for *p* = 0.10, 0.50 and 0.90, where predicted injury reaches 10, 50 and 90 % of the asymptote. Uncertainty was propagated by Monte Carlo sampling of 1,000 parameter sets from the multivariate normal distribution of the fitted parameters; draws with a negative rate or an asymptote outside the bounds were discarded (461 to 1,000 retained per curve), curves were clipped to [0, 3] and the 2.5th and 97.5th percentiles gave 95 % confidence intervals. A curve whose asymptote fell below 2.5 had not saturated; its LT_50_ is marked as extrapolated in Figure 2 and Table 1.

**Figure 2.**
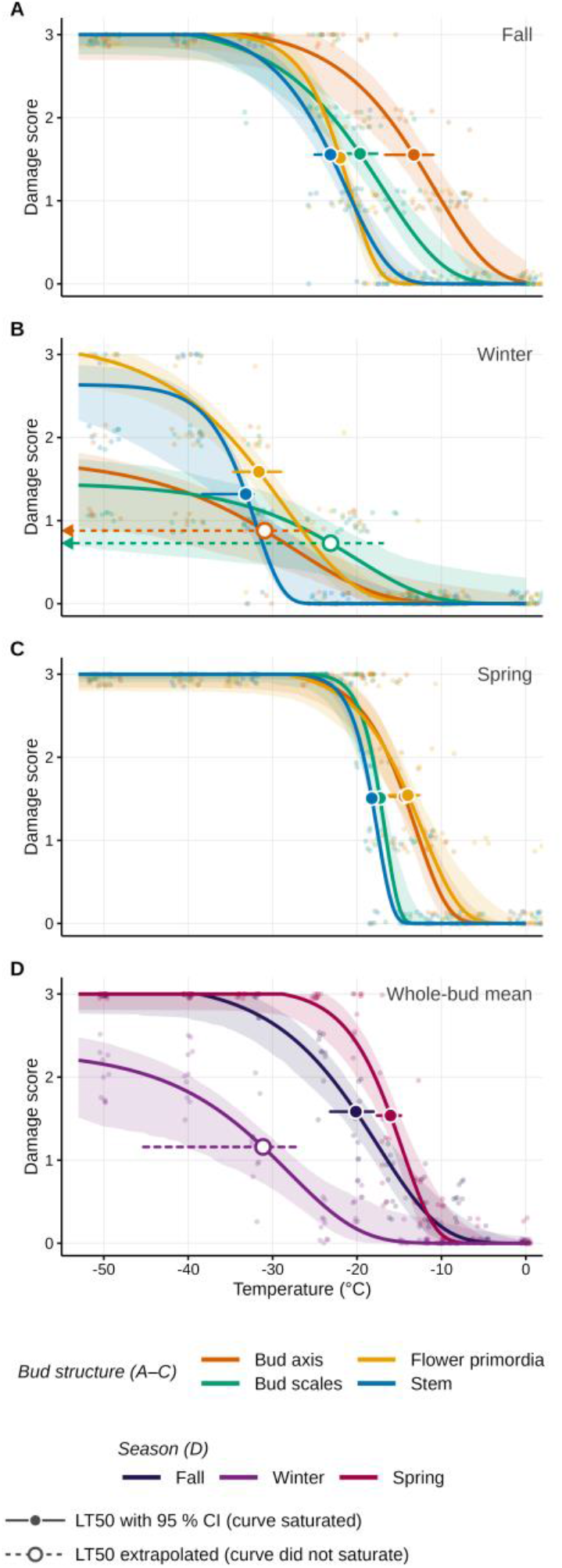
Freezing injury of *Vaccinium corymbosum* floral-bud structures in controlled freezing tests. Mean injury score (0 to 3) of the bud axis, bud scales, flower primordia and subtending stem against treatment temperature in (A) fall, (B) winter and (C) spring, and (D) the whole-bud mean by season. Faded circles are individual buds (*n* = 10 per temperature); lines are Gompertz fits with 95 % Monte Carlo confidence bands. Markers give LT_50_ with its 95 % confidence interval: filled marker and solid bar, saturated curve (asymptote ≥ 2.5); open marker and dashed bar, extrapolated LT_50_; arrowheads mark intervals extending beyond the plotted range.

**Table 1.** Seasonal summary of floral-bud traits of *Vaccinium corymbosum* measured on buds collected in fall 2021, winter 2022 and spring 2022. Fig., figure in which the variable is shown. KW, Kruskal-Wallis test. Letters after water-content means denote seasons that differ (pairwise Wilcoxon tests, Holm correction, *P* < 0.05).

| Variable | Fig. | Fall 2021 | Winter 2022 | Spring 2022 | Season effect |
| --- | --- | --- | --- | --- | --- |
| Whole-bud LT<sub>50</sub> (°C) [95 % CI]<sup>a</sup> | 1, 2 | -20.1 [-23.2, -18.0] | -31.1 [-45.3, -27.2]<sup>b</sup> | -16.0 [-17.6, -14.8] | all seasons differ (CI non-overlap) |
| Flower-primordium LT<sub>50</sub> (°C) [95 % CI]<sup>a</sup> | 2 | -22.0 [-23.1, -21.2] | -31.6 [-34.6, -29.0] | -14.0 [-16.2, -12.6] | all seasons differ |
| Bud water content (% fresh mass), mean ± SD, $n = 20$ | 1 | 48.2 ± 8.3 a | 49.8 ± 3.5 a | 59.1 ± 0.7 b | Kruskal-Wallis $\chi^2 = 35.5$, $P < 0.001$ |
| Stem nucleation temperature, IDTA (°C)<sup>c</sup> | 3 | n.d. | -13.0 | -11.8 |  |
| Delay from stem nucleation to bud freezing, IDTA (min), bud base / apex<sup>c</sup> | 3 | n.d. | 2.5 / 5.4 | 2.0 / 2.8 |  |
| Low-temperature exotherm, DTA (modules with LTE / modules run) | 3 | 0 / 9 | 0 / 8 | 0 / 4 | none in any season |
| Buds with frozen flower primordia at -10 °C | 4 | 0 of 5 | 0 of 5 | 5 of 5 |  |
| Xylem trace present (floret × landmark records) | 5, 6 | 12 of 12 | 12 of 12 | 12 of 12 | no change |
| Mature-type share of readable trace elements, rachis, mean ± SD, $n = 3$ | 6 | 0.07 ± 0.12 | 0.22 ± 0.19 | 0.73 ± 0.24 | exact KW $H = 6.21$, $P = 0.021$ |
| Mature-type share, pedicels of florets 1 to 3 pooled, mean ± SD, $n = 3$ | 6 | 0.06 ± 0.10 | 0.18 ± 0.16 | 0.56 ± 0.31 | exact KW $H = 4.51$, $P = 0.11$ |
| Conduit density, bud base (mm<sup>-2</sup>), mean ± SD, $n = 3$ | 7 | 9,663 ± 1,923 | 12,794 ± 293 | 11,276 ± 827 | exact KW $H = 5.96$, $P = 0.025$ |
| $D_h$, bud base (μm), mean ± SD, $n = 3$ | 7 | 2.53 ± 0.15 | 2.17 ± 0.31 | 2.47 ± 0.19 | exact KW $H = 2.76$, $P = 0.30$ |

<sup>a</sup> LT<sub>50</sub> from Gompertz fits to mean injury per temperature, with 95 % Monte Carlo confidence intervals (1,000 parameter draws). <sup>b</sup> Extrapolated: the whole-bud winter curve did not saturate within the tested range (fitted asymptote 2.32 on the 0 to 3 scale). <sup>c</sup> Values read from the representative winter and spring runs shown in Figure 3 (three buds per season behaved alike); n.d., not determined (camera not available at the fall collection). $D_h$ , weighted mean conduit diameter, $\Sigma d^4 / \Sigma d^3$ (see Methods).

### Thermal detection of freezing: infrared and differential thermal analysis

Ice nucleation and propagation were followed by IDTA, in which successive infrared images are subtracted so that the latent heat of freezing appears as a transient signal (Hacker & Neuner 2007; Wisniewski et al. 2015), on three buds each in winter and spring; the camera was not available at the fall collection. Stem segments bearing one floral bud stood with the base in moist potting medium, which seeds ice, in the Tenney Junior chamber, were brought to -5 °C within 1 h, held for 4 h and then cooled at 6 °C h⁻¹ until the stem froze, while recorded with an infrared camera (SC8303 with a 1x microscope close-up lens, 3 to 5 µm, f/4, thermal sensitivity below 25 mK; FLIR Systems, Wilsonville, OR, USA). The times at which freezing began in the stem, bud base and bud apex were read from the difference images in FLIR Research Studio (FLIR Systems), with the stem surface temperature at the first event taken as the nucleation temperature (Figure 3A, B).

**Figure 3.**
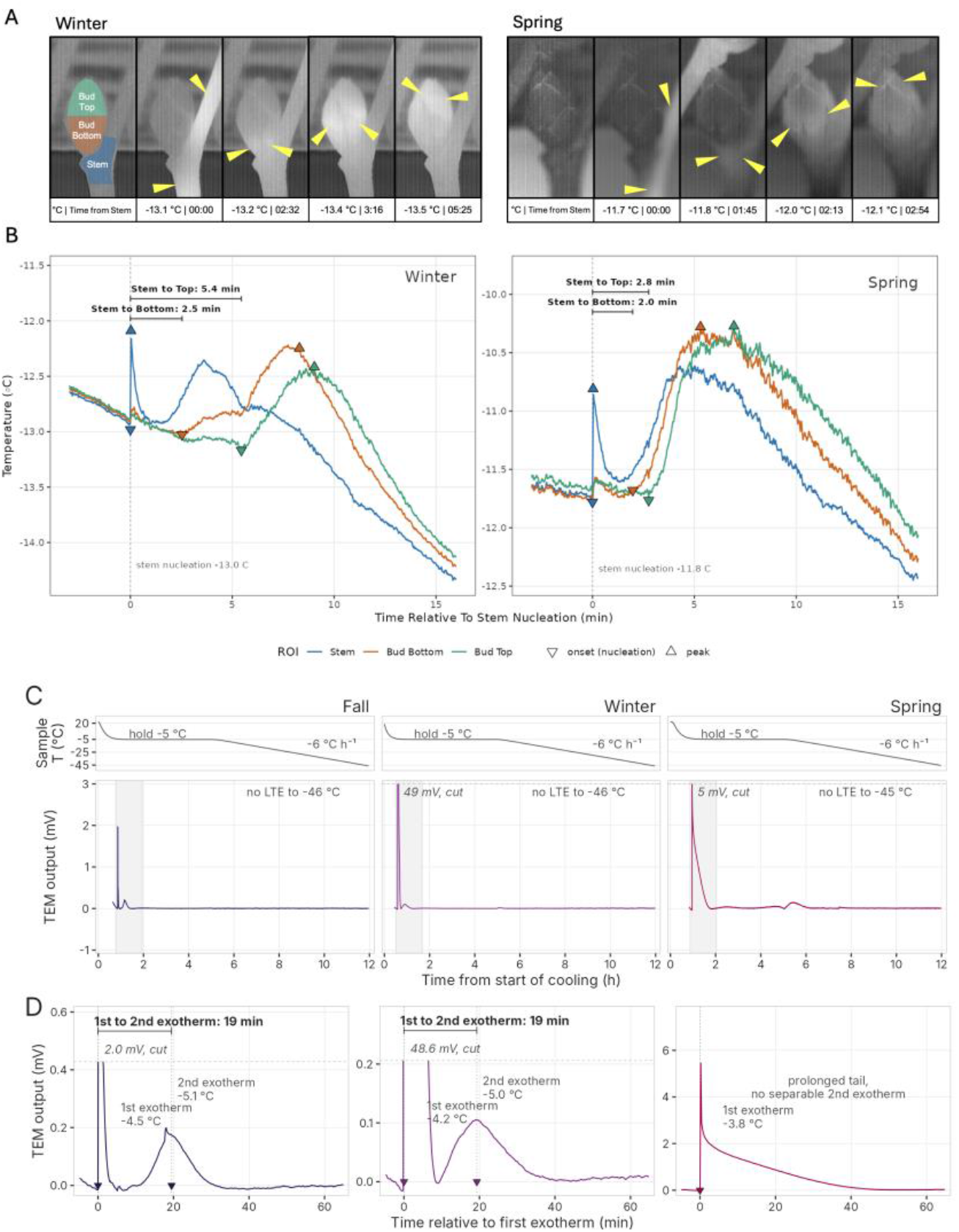
Thermal detection of freezing in *Vaccinium corymbosum* floral buds. (A) Infrared images of one of the three buds run per season, with attached stem and the stem base standing in moist medium, in winter (left) and spring (right); the first frame shows the regions of interest (stem, bud base, bud apex), each later frame gives the stem temperature and the time from stem nucleation, and yellow arrowheads mark the advancing freezing front. (B) Temperature of the three regions in the same runs against time from stem nucleation; downward triangles mark freezing onsets, upward triangles the peaks, and brackets the delays from stem nucleation to onset in the bud base and apex. (C) Differential thermal analysis of one module (two buds) per season: sample temperature (upper strips) and baseline-corrected output of the thermoelectric module (TEM; lower panels) from the start of cooling through the -5 °C hold and the ramp at 6 °C h⁻¹. First exotherms are clipped at the dashed line, with their peak values given; the grey band marks the window enlarged in D; no low-temperature exotherm (LTE) appeared during the ramp; the small bump between 4 and 6 h in spring (< 0.1 mV), present in every spring module, is a run artefact, not an LTE. (D) The hour after the first exotherm in the same modules: in fall and winter a sharp first exotherm is followed 19 min later by a broad second exotherm, the buds freezing slowly; in spring a single exotherm decays over a prolonged tail. Exotherm temperatures are given on the panels.

DTA used a multi-module system (Mills et al. 2006) in which the voltage of a thermoelectric module (HP-127-1.4-1.5-74, TE Technology, Traverse City, MI, USA) under each sample is logged against a reference by a multimeter data acquisition system (2700-DAQ-40, Keithley Instruments, Cleveland, OH, USA), with tray temperature read by a thermistor and chamber temperature by two type T thermocouples on a separate logger (RDXL4SD, Omega, Stamford, CT, USA). Buds were run within 24 h, two buds per module, each with about 5 mm of stem, under a square of moist paper towel that seeds nucleation. Nine modules were run in fall, nine in winter and nine in spring, 27 in all (42 buds; examples in Figure 3C, D). The chamber ran the same programme as for IDTA, the 4 h hold at -5 °C letting the first freezing events run to completion before the ramp towards -55 °C; thermograms were baseline-corrected with an asymmetrically weighted Whittaker smoother (Eilers 2003) and analysed to -46 °C.

### Cryomicroscopy of ice distribution

Ice was located by reflected-polarised-light cryomicroscopy (Stegner et al. 2020) on a bench inside a walk-in growth chamber (LTRB-37, BioChambers, Winnipeg, Canada), where a stereomicroscope (Stemi 2000-C, Zeiss, Jena, Germany), cold-light source (KL 1500 LCD, Zeiss), crossed polarising filters and a digital camera (D5100, Nikon, Tokyo, Japan) stood with all tools at chamber temperature; buds were handled only with tools so that hand heat did not warm them. Five buds per season, each with about 5 mm of stem, were sealed in bags with moist paper. The chamber, with the equipment inside, was cooled from room temperature (20 °C) at 2 °C h⁻¹ so that no condensation formed on the optics; the buds were introduced from 4 °C storage when the chamber reached 4 °C, cooled to 0 °C at the same rate and from 0 to -10 °C at 1 °C h⁻¹, and held at -10 °C for 9 h, above every seasonal LT_50_ but low enough for freezing to run to completion, to compare ice location in frozen but not lethally injured buds (Figure 4). At -10 °C each bud was sectioned longitudinally and transversely and photographed at once, ice appearing bright between crossed polarisers. Five zones (flower primordia, inner scales, outer scales, bud axis, subtending stem) were scored per bud from the photographs as holding a large ice mass, a small one or none, and as frozen or unfrozen from the appearance of the tissue.

**Figure 4.**
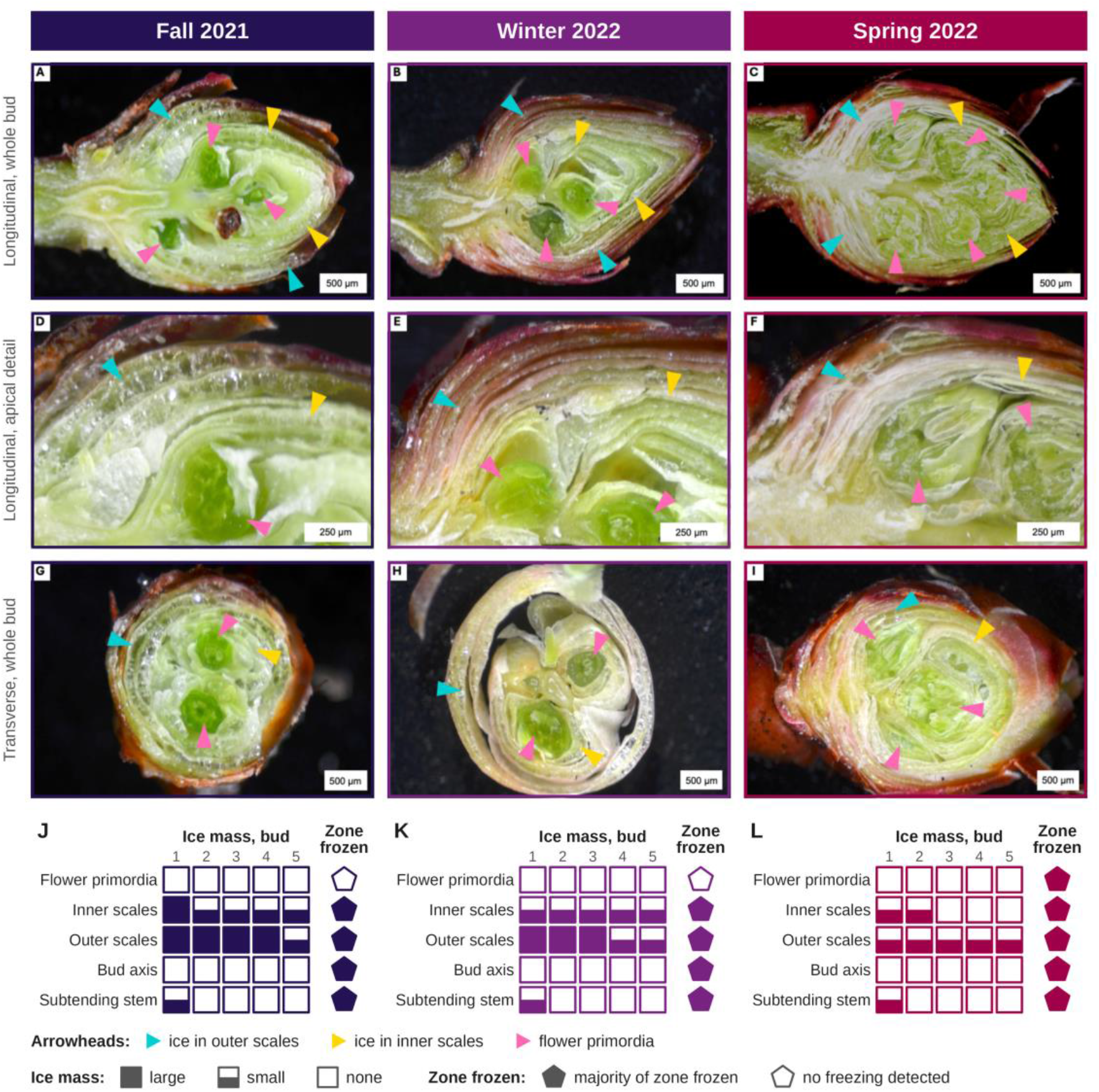
Ice distribution in five *Vaccinium corymbosum* floral buds after 9 h at -10 °C, by reflected-polarised-light cryomicroscopy. Representative buds in fall 2021 (A, D, G), winter 2022 (B, E, H) and spring 2022 (C, F, I): whole-bud longitudinal sections (A to C), apical detail of the longitudinal sections centred on the flower primordia (D to F) and whole-bud transverse sections (G to I). Ice appears bright; cyan and yellow arrowheads mark ice in the outer and inner scales, and pink arrowheads mark the flower primordia. Scale bars 500 µm (A to C, G to I) and 250 µm (D to F). (J to L) Ice in five zones of the five buds per season: filled square, large ice mass; lower half filled, small ice mass; empty square, none. Pentagons give one call per zone and season: filled, the majority of the zone frozen; open, no freezing detected.

### Light microscopy of the xylem trace to the flower primordia

Three buds per season, each with about 5 mm of stem, were fixed in 50 % ethanol, 5 % glacial acetic acid and 10 % formalin (33 % formaldehyde) in water, under vacuum for 1 h and at 4 °C overnight, dehydrated through an ethanol series (30 to 100 %), cleared in Histo-Clear II and embedded in paraffin (Paraplast X-tra) at 58 °C. Median longitudinal sections 6 µm thick were cut on a rotary microtome (Spencer Lens Co., Buffalo, NY, USA) with HP35 Ultra blades (Epredia, Kalamazoo, MI, USA), stained with safranin O, chlorazol black and fast green in ethanol, mounted in Permount and viewed on an Axio Imager.A2 microscope (Carl Zeiss, Jena, Germany) by epifluorescence, under which safranin-stained lignified walls fluoresce brightly (Bond et al. 2008), and by brightfield. One median section per bud was selected before scoring on the criterion that the rachis and the pedicels of the three most basal florets were cut longitudinally. Landmarks used throughout were the bud-stem union, the rachis immediately below the first (most basal) floret (R), the pedicels of florets 1 to 3 counted from the base (P1 to P3) and the receptacle base of each floret (Figure 5 labels the rachis and pedicels and marks the transverse-section planes). Xylem was recorded as present at a landmark when at least one tracheary element with secondary thickenings could be followed in longitudinal view (36 records in nine buds).

**Figure 5.**
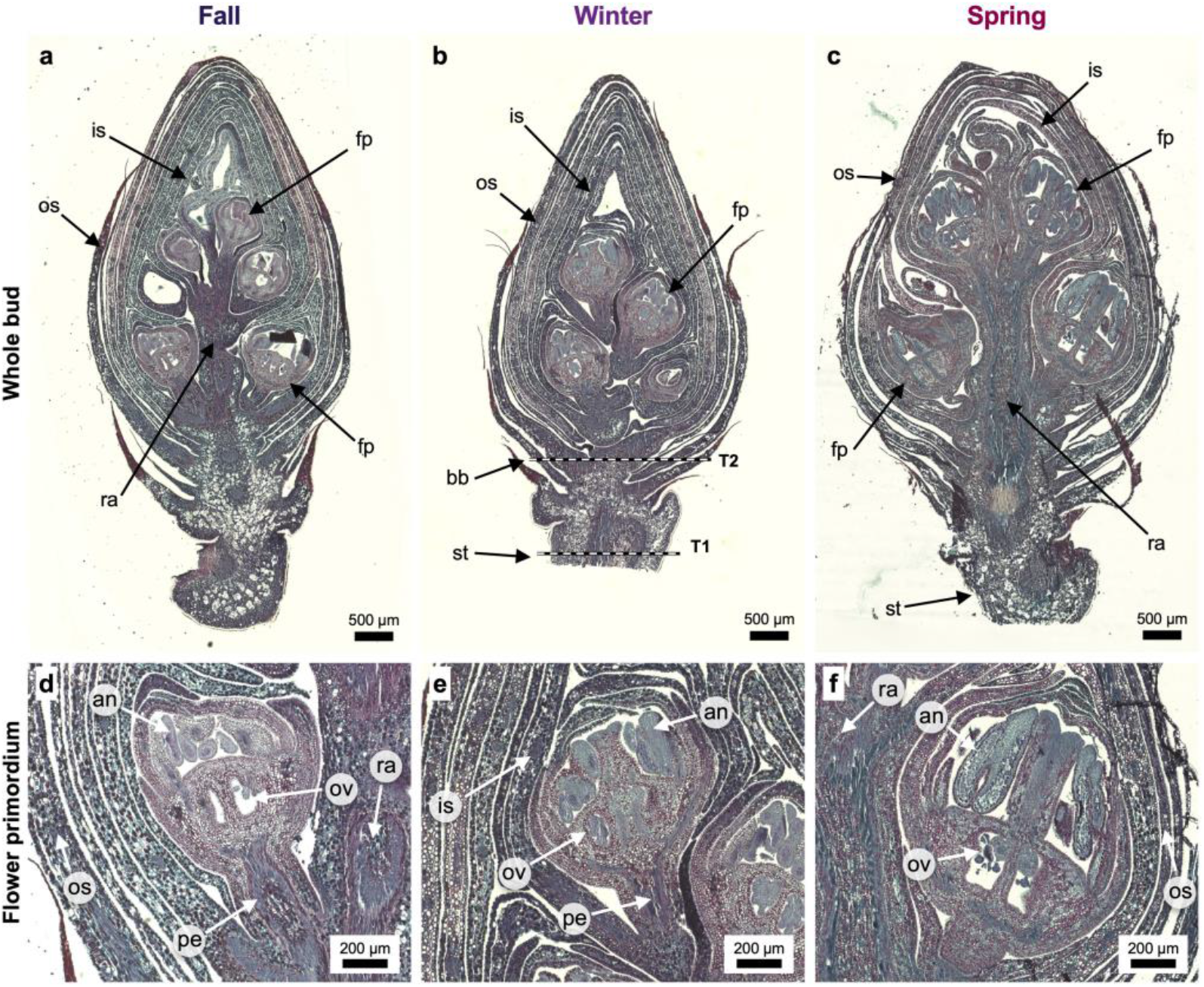
Anatomy of *Vaccinium corymbosum* floral buds in fall (A, D), winter (B, E) and spring (C, F), in median longitudinal paraffin sections stained with safranin O, chlorazol black and fast green and viewed in brightfield. (A to C) Whole buds with the subtending stem, one bud per season, at one magnification. (D to F) Flower primordia at higher magnification: E and F are basal primordia of the sections in B and C, D is from another fall section, and F is shown at lower magnification than D and E so that the whole primordium is included. Dashed lines in B are illustrative and show where the transverse sections of Figure 7, cut from other buds, were taken: T2, bud base immediately above the bud-stem union; T1, stem about 1 mm below the union. an, anthers; bb, bud base; fp, flower primordium; is, inner scales; os, outer scales; ov, ovary; pe, pedicel; ra, rachis; st, stem. Scale bars 500 µm (A to C) and 200 µm (D to F).

### Wall-pattern scoring of trace maturation

In the same sections, every tracheary element in longitudinal view at R and P1 to P3 was assigned to one of two classes (examples in Figure 6A to F): Type A, immature, with annular or helical thickenings separated by open primary wall, the protoxylem-type patterns of elongating tissue; Type B, mature, with a secondary wall covering most of the wall surface and leaving slits or pits, the scalariform, reticulate or pitted patterns of metaxylem, formed after elongation ceases (Evert 2006). Elements cut transversely or too obliquely to read were counted as seen but not classified, as were doubtful elements. Every call was made at the microscope on a single focal plane, at 40x and, for unresolved elements, at 63x. Because half of the pedicel landmarks held two or fewer readable elements, counts were summed over the three pedicels of each bud, the rachis kept its single landmark, and the response was the proportion of mature-type elements among classified elements, B/(A + B), with the bud as the unit of replication (*n* = 3 per season; Table 2). Figures 5 and 6 combine mosaics and extended-focus composites, adjusted only globally (Cromey 2010), with background outside the original frames filled and marker ink masked, using scripts in Python with NumPy and SciPy (Harris et al. 2020; Virtanen et al. 2020); composites were not used for scoring.

**Figure 6.**
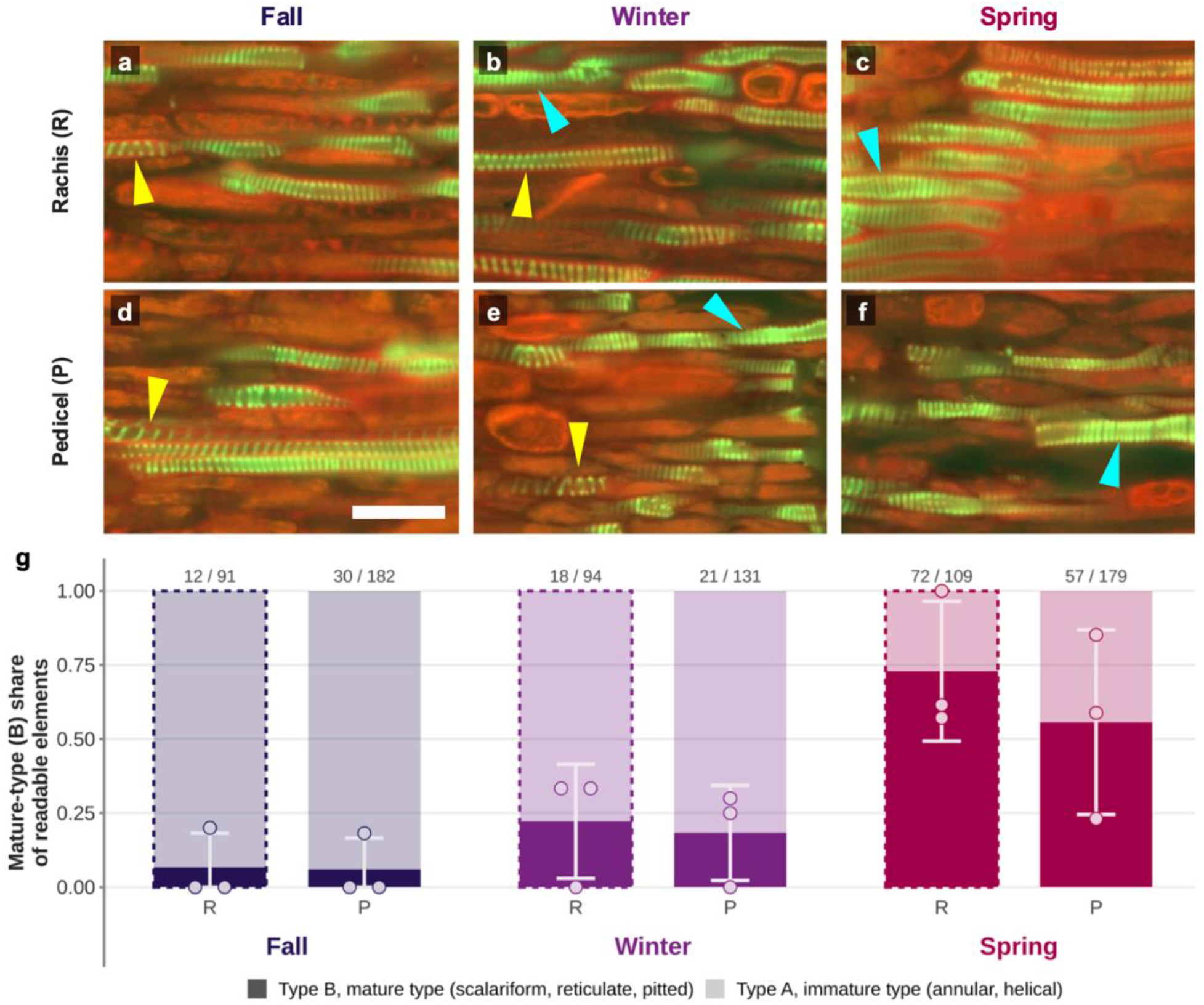
Maturation of the xylem trace to the flower primordia of *Vaccinium corymbosum* floral buds. (A to F) Tracheary elements in median longitudinal paraffin sections stained with safranin O, chlorazol black and fast green and viewed by epifluorescence, at the rachis immediately below the first floret (A to C) and in the pedicel of floret (D to F), in fall (A, D), winter (B, E) and spring (C, F); each panel is from one bud. Yellow arrowheads, helical elements (Type A, immature: annular or helical thickenings separated by open primary wall); cyan arrowheads, scalariform elements (Type B, mature: secondary wall covering most of the wall surface and leaving narrow slits). B, C, E and F are extended-focus composites of four, three, four and two focal planes, A and D single planes; elements were scored on single planes. Scale bar is 20 µm. (G) Composition of the readable tracheary elements by season at the rachis (R, dashed outline) and in the pedicels of florets 1 to 3 pooled (P): bars, mean of three buds divided into mature-type (solid) and immature-type (tint) shares; circles, individual buds (mature-type share); whiskers, standard deviation of the bud values clipped at 0 and 1; numbers above the bars, elements read / elements seen.

**Table 2.** Maturation state of the xylem trace in *Vaccinium corymbosum* floral buds by season, from one median longitudinal section per bud (*n* = 3 buds per season). Tracheary elements seen, elements observed at the landmark; classified, elements in longitudinal view assigned to a wall-pattern class (Type A, annular or helical, immature; Type B, scalariform, reticulate or pitted, mature); mature-type, Type B elements. Counts are summed over the three buds of each season; pedicel counts are also summed over florets 1 to 3, and the rachis was scored once per bud, immediately below floret 1. Mature-type share, B/(A + B) per bud, mean ± SD of the three buds.

| Landmark | Season | Tracheary elements seen | Classified (Type A + B) | Mature-type (Type B) | Mature-type share, mean ± SD |
| --- | --- | --- | --- | --- | --- |
| Rachis below floret 1 | Fall | 91 | 12 | 1 | $0.07 \pm 0.12$ |
| | Winter | 94 | 18 | 4 | $0.22 \pm 0.19$ |
| | Spring | 109 | 72 | 52 | $0.73 \pm 0.24$ |
| Pedicels of florets 1 to 3 | Fall | 182 | 30 | 2 | $0.06 \pm 0.10$ |
| | Winter | 131 | 21 | 4 | $0.18 \pm 0.16$ |
| | Spring | 179 | 57 | 36 | $0.56 \pm 0.31$ |

Xylem was present at every landmark in every bud (36 of 36 records). Season effect on share, bud as unit, exact permutation Kruskal-Wallis (1,680 label assignments): rachis $H = 6.21$ , $P = 0.021$ ; pedicels $H = 4.51$ , $P = 0.11$ . Hedges' $g$ [exact 95 % CI], later season minus earlier: rachis, spring vs fall 2.85 [0.53, 5.11], spring vs winter 1.88 [0.04, 3.63], winter vs fall 0.78 [-0.64, 2.13]; pedicels, spring vs fall 1.71 [-0.06, 3.37], spring vs winter 1.21 [-0.36, 2.67], winter vs fall 0.72 [-0.68, 2.05]. Exact pairwise permutation $P = 0.10$ (the smallest attainable at 3 vs 3) for spring vs fall at both landmarks.

### Conduit anatomy at the bud-stem junction

For conduit anatomy, buds with about 5 mm of stem were fixed in 4 % glutaraldehyde overnight at 4 °C, rinsed in 0.05 M potassium phosphate buffer, dehydrated in ethanol (30 to 100 %) and embedded in medium-grade LR White resin (Ted Pella, Redding, CA, USA) cured at 60 °C for 28 h (Villouta et al. 2022). Transverse sections 2 µm thick were cut on an ultramicrotome (MT-2, Sorvall, Norwalk, CT, USA) from three buds per season at two planes (Figure 5B): the bud base immediately above the bud-stem union (T2) and the subtending stem about 1 mm below it (T1). Sections were stained with methylene blue and azure II, then basic fuchsin, and photographed in brightfield on the same Axio Imager.A2 microscope. In ImageJ (Schneider et al. 2012), every conduit within a 20,000 µm² pie-sliced sector of each section was counted and its lumen outlined; the wall thickness (*t*, a single wall, not the double wall between neighbouring lumens) and the lumen span (*b*) were measured on ten randomly selected conduits, and the stele and the xylem ring were outlined on the whole section.

Six traits were derived (Figure 7; Table 3) (Scholz et al. 2013): conduit density, the number of conduits per mm² of sector; a weighted mean conduit diameter, *D*ₕ = Σ*d*⁴/Σ*d*³, where *d* is the diameter of a circle with the area of each lumen, which gives wide conduits less weight than the conductivity-weighted Σ*d*⁵/Σ*d*⁴ (Sperry et al. 1994); the implosion-resistance index (*t*/*b*)², from the section means of *t* and *b*, which with a single-wall *t* is one quarter of the double-wall index of Hacke et al. (2001); lumen area fraction, total lumen area over sector area; xylem fraction of the stele, xylem-ring area over stele area; and stele cross-sectional area. Two further descriptors were calculated: the diameter of a circle with the mean lumen area of a section (lumen area fraction divided by conduit density), and the non-xylem area of the stele (stele area minus xylem-ring area).

**Figure 7.**
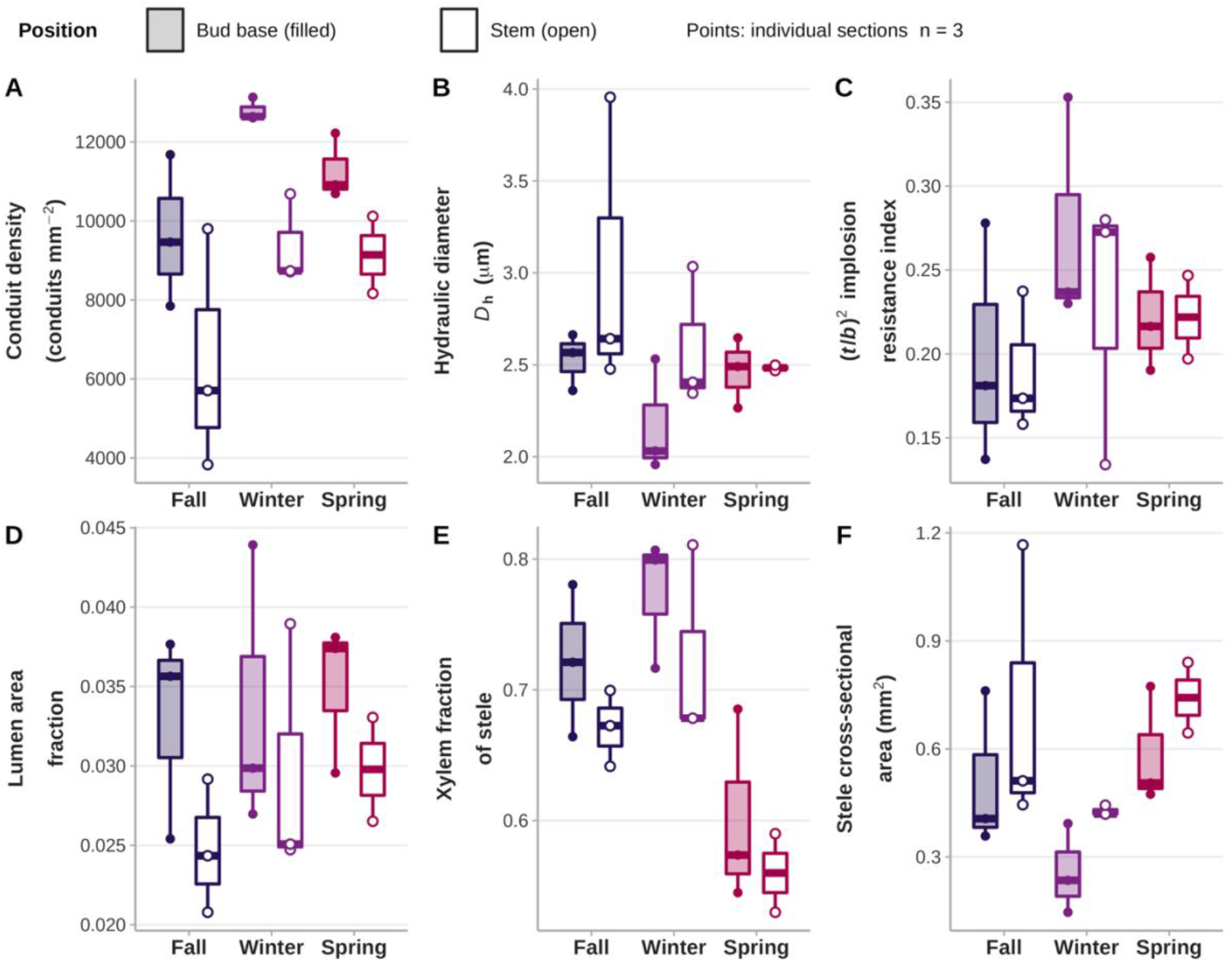
Conduit anatomy at the bud-stem junction of *Vaccinium corymbosum* floral buds across the dormancy cycle. Traits of transverse sections through the bud base immediately above the bud-stem union (plane T2 in Figure 5B; filled boxes) and through the subtending stem about 1 mm below it (plane T1; open boxes) in fall 2021, winter 2022 and spring 2022. (A to D) Traits measured within a sector of the xylem ring: (A) conduit density, (B) weighted mean conduit diameter *D*ₕ, (C) implosion-resistance index (*t*/*b*)² and (D) lumen area fraction. (E, F) Traits measured on the whole section: (E) xylem fraction of the stele and (F) stele cross-sectional area. Boxes show the median and interquartile range, whiskers extend to the most extreme value within 1.5 times the interquartile range, and points are individual sections (*n* = 3 per season and position). Season colours as in all figures.

**Table 3.** Season effects on conduit and tissue traits at the bud base (plane T2) and in the subtending stem (plane T1) of *Vaccinium corymbosum* floral buds; the section values are shown in Figure 7. Exact permutation Kruskal-Wallis *H* with *P* for the season effect within position (1,680 label assignments for 3/3/3; 560 for 3/3/2), and Hedges’ *g* with exact 95 % confidence intervals for the three seasonal contrasts (later season minus earlier; positive, higher in the later season); *n* = 3 sections per group.

| Trait | Position | Exact KW $H$ ($P$) | Winter vs fall $g$ [95 % CI] | Spring vs winter $g$ [95 % CI] | Spring vs fall $g$ [95 % CI] |
| --- | --- | --- | --- | --- | --- |
| Conduit density | Bud base | 5.96 (0.025) | 1.82 [0.00, 3.54] | -1.96 [-3.74, -0.08] | 0.87 [-0.58, 2.24] |
|  | Stem | 1.89 (0.446) | 1.02 [-0.48, 2.42] | -0.14 [-1.44, 1.17] | 0.75 [-0.72, 2.12] |
| $D_h$ | Bud base | 2.76 (0.296) | -1.16 [-2.60, 0.39] | 0.91 [-0.55, 2.28] | -0.29 [-1.56, 1.03] |
|  | Stem | 1.89 (0.446) | -0.54 [-1.84, 0.82] | -0.26 [-1.56, 1.08] | -0.60 [-1.93, 0.82] |
| $(t/b)^2$ | Bud base | 1.87 (0.439) | 0.85 [-0.60, 2.20] | -0.76 [-2.10, 0.65] | 0.32 [-0.99, 1.60] |
|  | Stem | 1.11 (0.671) | 0.48 [-0.87, 1.77] | -0.07 [-1.37, 1.24] | 0.59 [-0.83, 1.92] |
| Lumen area fraction | Bud base | 0.36 (0.879) | 0.07 [-1.22, 1.35] | 0.16 [-1.14, 1.43] | 0.30 [-1.02, 1.57] |
|  | Stem | 2.00 (0.425) | 0.60 [-0.78, 1.91] | 0.02 [-1.28, 1.32] | 0.84 [-0.66, 2.24] |
| Xylem fraction of stele | Bud base | 5.07 (0.086) | 0.77 [-0.65, 2.11] | -2.19 [-4.08, -0.20] | -1.45 [-3.00, 0.21] |
|  | Stem | 4.75 (0.071) | 0.71 [-0.69, 2.04] | -1.76 [-3.57, 0.16] | -2.37 [-4.55, -0.11] |
| Stele cross-sectional area | Bud base | 5.07 (0.086) | -1.12 [-2.55, 0.41] | 1.79 [-0.01, 3.49] | 0.31 [-1.00, 1.59] |
|  | Stem | 5.20 (0.061) | -0.79 [-2.14, 0.63] | 2.83 [0.30, 5.31] | 0.08 [-1.23, 1.37] |

$D_h$ , weighted mean conduit diameter, $\Sigma d^4/\Sigma d^3$ (see Methods); $(t/b)^2$ , implosion-resistance index with single-wall $t$ (one quarter of the double-wall form).

### Statistical analyses

Water content was compared among seasons by the Kruskal-Wallis test (fall and winter distributions were not normal) with pairwise Wilcoxon tests and Holm correction; lethal temperatures by non-overlap of 95 % confidence intervals. The anatomical datasets have three buds per season, the usual replication for this kind of work (Kuprian et al. 2017; Neuner et al. 2019b), so all tests on them are exact: the season effect within each landmark or position was tested by a permutation Kruskal-Wallis test over all assignments of buds to seasons, and pairwise contrasts were summarised by bias-corrected Hedges’ *g* (Hedges 1981) with exact 95 % confidence intervals from the noncentral *t* distribution (Cumming & Finch 2001). Within buds, the rachis was compared with the pooled pedicels, and floret 1 with floret 3, by exact sign-flip tests on the paired differences, and the rachis-pedicel contrast at the element level by a Cochran-Mantel-Haenszel odds ratio stratified by bud with a permutation *P*. Whether the binary ‘Type B present’ score depended on the number of readable elements was tested by an exact Mann-Whitney *U* test. Covariation of each conduit trait with stele area was tested by Spearman rank correlation across sections and within each position. Analyses used R 4.3.3 (R Core Team 2023) with ggplot2 3.4.4 (Wickham 2016) and minpack.lm 1.2-4.

## Results

### Bud water content and whole-bud hardiness

Mean bud water content was the same in fall and winter and rose in spring (Figure 1A; Table 1): 48.2 ± 8.3, 49.8 ± 3.5 and 59.1 ± 0.7 % of fresh mass (Kruskal-Wallis χ² = 35.5, df = 2, *P* < 0.001; Holm-corrected *P* = 0.80 for fall versus winter and *P* < 0.001 for spring versus either). The spread among buds, however, narrowed at each step (coefficient of variation 17, 7 and 1 %). The fall sample was bimodal (six buds at 35.6 to 39.0 %, drier than any winter bud; eleven at 53.6 to 56.6 %; three intermediate), so its mean describes a mixture of hydration states; by winter the buds had converged on one state (18 of 20 between 46.1 and 50.2 %), and spring buds more so.

Whole-bud LT_50_ from the controlled freezing tests followed the expected cycle (Figure 1B; Table 1): - 20.1 °C (95 % CI -23.2 to -18.0) in fall, -31.1 °C (-45.3 to -27.2) in winter and -16.0 °C (-17.6 to -14.8) in spring, the three intervals non-overlapping. Buds therefore gained 11.0 °C of hardiness between fall and winter while mean water content stayed the same and the spread among buds narrowed (CV 17 to 7 %), and lost 15.1 °C between winter and spring while water content rose by about 9 percentage points. The wide winter interval reflects a whole-bud injury curve that had not saturated at the lowest test temperature (asymptote 2.32 on the 0 to 3 scale), so the winter LT_50_ is partly extrapolated; its upper limit still lies below the fall interval.

### Freezing injury by bud structure

Gompertz curves fitted every structure in every season (Figure 2). In fall (Figure 2A) the stem and the flower primordia were the hardiest structures (LT_50_ -23.2 and -22.0 °C), the scales intermediate (-19.6 °C) and the bud axis by far the most sensitive (-13.3 °C, 95 % CI -16.6 to -11.0; LT₁₀ -5.2 °C), resolved from every other structure. In winter (Figure 2B) every structure was hardier; the axis and scale curves did not saturate (asymptotes 1.76 and 1.46), so their LT_50_ values (-30.9 and -23.1 °C) are extrapolated with wide intervals, the primordia (-31.6 °C, -34.6 to -29.0) and the stem (-33.2 °C, -38.3 to -32.3) saturated, and no pair of winter structures was resolved. In spring (Figure 2C) the order inverted: the stem (-18.2 °C) and the scales (-17.3 °C) were hardier than the primordia (-14.0 °C, -16.2 to -12.6) and the axis (-14.4 °C, - 16.1 to -13.2), which showed measurable injury (LT₁₀) at -8.5 and -9.9 °C. Flower-primordium LT_50_ thus deepened by 9.6 °C from fall to winter and rose by 17.6 °C from winter to spring, a larger spring loss than the whole bud’s 15.1 °C (Figure 2A to D; Table 1).

### Thermal analysis of freezing

IDTA showed the same order of events in all three buds per season in winter and spring (Figure 3A, B), stem first, then bud base, then apex, but at a different pace between seasons. The stem nucleated at -13.0 and -11.8 °C in the runs shown (-11.8 to -14.5 °C across runs) and froze abruptly; ice reached the bud base 2.5 and 2.0 min later and the apex after 5.4 and 2.8 min, and the bud froze slowly, over 8 to 11 min. In winter the bud thus froze as two separable events, the apex lagging the base by 2.9 min, with a low, drawn-out exotherm (about 0.8 °C above the cooling trend); in spring it froze as a single event, base and apex together (0.8 min apart), with a larger exotherm (about 1.4 °C). DTA on buds under moist paper reproduced the two-stage pattern (Figure 3C, D). All 21 modules produced a single exotherm complex at or near the -5 °C hold and nothing thereafter: during the ramp to -46 °C the largest deviation above baseline was 0.011 to 0.080 mV, below 2.5 % of the first exotherm in fall and winter and 10 % in spring, so no LTE was detected. In fall and winter the complex began with a sharp exotherm (onset -2.2 to -5.3 °C) from the free water of the paper, cut surface and stem, followed by a broad, low exotherm in seven of nine fall and three of eight winter modules, 12 to 22 min later in all but one (75 min in one fall module); the broad event has the duration and timing of the bud freezing seen by IDTA. In spring one module gave a sharp spike with a 60 min tail and three only slow exotherms. DTA nucleated at -2 to -5 °C, with ice on the paper around the bud; in IDTA, with only the moist medium at the twig base as an ice source, the stem stayed unfrozen through the 4 h hold at -5 °C and froze at -12 to -14 °C during the ramp.

### Ice distribution at -10 °C

After 9 h at -10 °C, ice masses were found in the scales in every season and in none of the flower primordia (Figure 4). In fall, large ice masses filled the outer scales of four of five buds and the inner scales of one, small masses occupied the remaining scale zones, and the primordia were ice-free and unfrozen in appearance (Figure 4A, D, G, J); in winter the pattern was the same with less ice (large masses in the outer scales of three buds, small masses in the inner scales of all five), the primordia again unfrozen (Figure 4B, E, H, K). In spring no large mass was seen, small masses persisted in the outer scales of all five buds and the inner scales of two, and the primordia, though without a discrete ice mass, were frozen in every bud, with the glassy, water-soaked appearance of frozen tissue (Figure 4C, F, I, L). Buds with frozen primordia thus rose from 0 of 5 in fall and winter to 5 of 5 in spring (Table 1). The bud axis held no discrete ice mass but was scored as frozen in every season (Figure 4J to L), so in fall and winter ice had passed the junction and the axis and was stopped only at the primordia; the subtending stem carried a small mass in one bud per season.

### Xylem trace to the flower primordia

In median longitudinal sections (Figure 5) a strand of tracheary elements ran from the stem through the bud-stem union, up the bud axis and rachis and into the pedicel of every floret examined, ending at the receptacle base. Xylem was present at all 36 floret by landmark records in the nine buds, so the continuity of the trace did not differ among seasons, and at the pedicel the trace was a single narrow strand a few elements wide in every season. This confirms in sections the continuous lignified strands that Flinn and Ashworth (1994b) saw in cleared buds, so any seasonal change in the vascular connection had to lie in the state of the elements rather than in their presence.

### Wall-pattern maturation of the trace

The differentiation state of the trace changed with season (Figure 6; Table 2). In fall the marked elements were helical, with open wall between the turns, and in the pedicel the turns were oblique and widely spaced (Figure 6A, D; yellow arrowheads). Winter fields showed the same loosely coiled helical elements next to occasional scalariform ones (Figure 6B, E). In spring the marked elements at both landmarks were scalariform, with narrow slits between closely spaced bars (Figure 6C, F; cyan arrowheads). The scores agreed with the images (Figure 6G). In fall and winter, mature-type elements were 0 to 0.33 of the readable elements of a bud at either landmark (season means 0.06 to 0.22), and three of the six dormant-season buds had none in any pedicel. In spring they were 0.57 to 1.00 at the rachis (mean 0.73) and 0.23 to 0.85 in the pedicels (mean 0.56).

The season effect was resolved at the rachis (exact permutation Kruskal-Wallis *H* = 6.21, *P* = 0.021) but not at the pedicels (*H* = 4.51, *P* = 0.11). Spring differed from fall with large effects at both landmarks (Hedges’ *g* = 2.85 at the rachis, 1.71 at the pedicels; exact pairwise *P* = 0.10, the smallest attainable), and from winter at the rachis (*g* = 1.88) but not detectably at the pedicels (1.21); fall and winter did not differ (Table 2). Excluding the six pedicel landmarks with a single readable element left the pattern unchanged (*H* = 4.75, *P* = 0.10). Within buds there was no front of maturation: the rachis was no more mature than the pooled pedicels of the same bud (paired difference +0.07; exact sign-flip *P* = 0.44; Cochran-Mantel-Haenszel odds ratio 1.34, *P* = 0.39), and floret 1 no more mature than floret 3 (difference -0.10; *P* = 0.59). Maturation in spring therefore appeared along the whole length of trace scored rather than advancing from the rachis into the pedicels.

Two features of the data bound these estimates. Only 210 of the 786 elements seen (27 %) could be read, half of the pedicel landmarks rested on two or fewer readable elements, and readability was lowest in the dormant seasons (16 % against 32 % in spring at the pedicels; 16 % against 66 % at the rachis), where the observer noted suspected mature-type patterns among unreadable elements in four buds; the fall and winter proportions are therefore lower bounds. A binary ‘Type B present’ score tracked sampling depth rather than season (recorded at 12 of 13 landmarks with six or more readable elements but at 5 of 23 with fewer; exact Mann-Whitney *P* = 0.0006), which is why the proportion among readable elements was used as the outcome.

### Conduit anatomy at the bud-stem junction

Conduits were narrow in every season at both positions, the bud base just above the bud-stem union (plane T2) and the subtending stem about 1 mm below it (plane T1) (Figures 5B, 7B). Group means of *D*ₕ lay between 2.2 and 3.0 µm, single sections between 1.96 and 3.96 µm, and unweighted mean lumen diameter between 1.65 and 2.84 µm. The bud base carried more conduits per unit area than the stem in every season and narrower conduits in fall and winter, with equal *D*ₕ in spring (Figure 7A, B).

At the bud base the conduit traits shifted in winter and returned towards the fall state in spring (Figure 7A to C; Table 3). Conduit density rose by 32 % from fall to winter and fell by 12 % from winter to spring, and the three winter sections exceeded every fall and spring section; the only conduit-trait season effect resolved (*H* = 5.96, *P* = 0.025; effect sizes in Table 3). Over the same interval *D*ₕ fell by 14 % and (*t*/*b*)² rose by 38 %, moderate to large effects that were not resolved (winter minus fall *g* = -1.16 and 0.85; season effect *P* = 0.30 and 0.44). The stem did not follow: conduit density rose by 45 % from fall to winter and did not reverse in spring, against a coefficient of variation of 47 % among fall sections, and no stem trait showed a season effect (*P* ≥ 0.42).

Lumen area fraction did not change with season at either position (*P* = 0.88 at the bud base, 0.43 in the stem; Figure 7D), so the share of the sector occupied by lumen stayed constant while conduit number and diameter changed. Xylem fraction of the stele was similar in fall and winter and lower in spring at both positions (Figure 7E), by 22 % from winter at the bud base and by 17 % from fall in the stem, the only contrasts of this trait whose intervals excluded zero (season effect *P* = 0.086 and 0.071).

Section size varied with season (Figure 7F): stele cross-sectional area was smallest in winter at both positions, 0.51 ± 0.22, 0.26 ± 0.13 and 0.58 ± 0.17 mm² (mean ± SD) at the bud base in fall, winter and spring (*H* = 5.07, *P* = 0.086), and 0.71 ± 0.40, 0.43 ± 0.01 and 0.74 ± 0.14 mm² in the stem (*H* = 5.20, *P* = 0.061). The xylem ring followed the same course (bud base 0.37, 0.20 and 0.35 mm²; stem 0.48, 0.31 and 0.41 mm²; *P* ≥ 0.23) and was smaller at the bud base than in the stem in every season. In spring the ring was as large as in fall, but the stele outside the ring was larger (bud base 0.24 against 0.14 mm², season effect *P* = 0.029; stem 0.33 against 0.23 mm², *P* = 0.19), so the lower spring xylem fraction came from tissue outside the ring, not from less xylem. Across the 17 sections, *D*ₕ increased with stele area (Spearman ρ = 0.54, *P* = 0.028; within the bud base ρ = 0.70, *P* = 0.043), whereas conduit density (ρ = - 0.51, *P* = 0.037) and xylem fraction (ρ = -0.62, *P* = 0.008) decreased with it, and lumen area fraction was unrelated to size (ρ = -0.12, *P* = 0.65). The winter shift of the bud-base conduit traits therefore coincided with the smallest sections.

## Discussion

The results met every expectation but one. As Flinn and Ashworth (1994b) found, hardiness peaked in winter and was lost in spring, ice formed in the scales while the primordia stayed ice-free in hardy buds, and a xylem trace reached the primordia in every season. The exception was the junction: winter bud-base sections had the expected denser, narrower conduits, but they were also the smallest sections, which alone could explain it. Read together (Table 1), the variables separate the two transitions. From fall to winter, LT_50_ deepened by 11 °C while ice exclusion and the state of the trace did not change. Mean water content did not change either, although its spread among buds narrowed. From winter to spring, LT_50_ rose by 15 °C and most variables changed: the trace matured, the primordia froze, water content rose and the xylem fraction of the stele fell. Only the order of freezing stayed the same, with nucleation in the stem, a delay of minutes before ice reached the bud, and no LTE. The one anatomical change established is the maturation of a trace that is present all year. We propose that it is part of the barrier Flinn and Ashworth left unexplained. What deepens hardiness from fall to winter is not resolved by the anatomy.

### Maturation, not continuity, tracks the loss of ice exclusion

Light microscopy, as in Flinn and Ashworth (1994b), rules out for *V. corymbosum* the simplest version of the *Prunus* model (Ashworth 1984; Ashworth et al. 1989; Quamme et al. 1995; Houghton et al. 2024): the dormant primordia are not vessel-free and no gap closes in spring. The wall-pattern data add the variable their stain could not resolve. In fall and winter, the readable elements of the trace were mostly annular and helical, the protoxylem type; in spring, scalariform, reticulate and pitted elements predominated (Figure 6). The dormant-season trace is therefore immature rather than absent, and the vessel proliferation they saw at deacclimation completes its differentiation. Nor does the species match *Calluna* and *Vitis*, where a mature connection is present and ice is stopped by a constriction or a permeability barrier (Jones et al. 2000; Kuprian et al. 2016), because the hardy *V. corymbosum* connection is not mature. Cranberry is the closer case: its bud-base tracheids were proposed to be not fully functional through winter (Villouta et al. 2022), a claim, like ours, about the state of a present connection. Our result also extends to a pre-formed trace the spring reconnection of bud and stem (Savage & Chuine 2021; McMann et al. 2022) and the vessel development that deacclimation induces within buds (Kasuga et al. 2023). Two features bear on mechanism. Maturation did not advance as a front from the rachis into the pedicels, so the switch is a property of the whole segment, not of a single choke point. And fall and winter were indistinguishable, so the anatomical change is confined to the interval in which exclusion is lost. Maturation was not all-or-none: at least a fifth of the readable winter elements (8 of 39) were already of the mature type (Figure 6B, E), yet primordia of the same collection stayed ice-free (Figure 4). A few mature elements do not let ice through; its entry may require most of the trace to mature, as in spring (88 of 129), or a second change acting with maturation, such as hydration of the trace (below).

The wall pattern marks developmental stage; it does not measure permeability to ice. Annular and helical elements conduct water in any elongating organ (Evert 2006), so the immature type does not by itself imply a closed pathway. Three properties of an arrested protoxylem-type strand could nonetheless make it a poor ice conductor. First, elements laid down during bud formation and then stretched as the bud enlarged may be partly occluded or collapsed. The loose, widely spaced coils in dormant-season pedicels (Figure 6D, E) are what protoxylem elements strained after deposition look like (Evert 2006). Second, the wall pattern does not tell whether the elements scored as immature had completed wall deposition and programmed cell death and were open conduits, and the openings that ice must cross, the pits and wall pores noted in the Introduction, are what matter (Ashworth & Abeles 1984; Wisniewski et al. 1991).

Third, dormant buds are poorly connected hydraulically (Savage & Chuine 2021), and a pathway that is not water-filled cannot conduct ice. Flinn and Ashworth (1994b) raised this possibility for the rachis and pedicels, and hydration alone removes supercooling from isolated peach primordia and raises the lethal temperature of *Vaccinium* buds (Bittenbender & Howell 1975; Rajashekar 1989), so rehydration and differentiation may open the gate together. The discriminating experiment is to follow conductivity and anatomy in one cohort across deacclimation: dye uptake (Signorelli et al. 2020) or imaging of bud water by magnetic resonance or X-ray phase contrast (Ishikawa et al. 1997, 2016; Kovaleski et al. 2019; Villouta et al. 2020) should show the trace becoming conductive and the primordia hydrating as exclusion is lost.

### What the junction contributes

Whatever the junction contributes to ice exclusion, it is not through the size of its conduits. First, they are too narrow to embolise on freezing and thawing: their *D*ₕ of 2 to 3 µm lies an order of magnitude below the 30 to 44 µm at which a single freeze-thaw cycle embolises conduits at moderate tension (Sperry & Sullivan 1992; Davis et al. 1999; Pittermann & Sperry 2003), and below the critical diameter of about 6 µm predicted even at -8 MPa (Pittermann & Sperry 2006). Second, lumen width cannot slow ice, because confinement depresses the freezing point only in pores of cell-wall and pit-membrane size, far narrower than a lumen (Ashworth & Abeles 1984; Wisniewski et al. 1991); a winter change in *D*ₕ of less than half a micrometre is therefore immaterial. The cranberry bud base, whose tracheids kept a conductivity-weighted diameter of 3 to 4 µm in every season (Villouta et al. 2022), gives the same picture. Third, the bud axis was frozen in every season, so ice crossed the junction whatever its anatomy, and the barrier lies above it, between axis and primordia (Figure 4).

The winter bud-base phenotype, denser and narrower conduits with a higher (*t*/*b*)², is confounded with section size. The xylem ring is lignified and its conducting elements are dead at maturity (Evert 2006), so the January ring should not be nearly half the size of the October one. Smaller buds, a cut farther above the union where the stele tapers, or a different outlining criterion are likelier explanations. Conduit diameter widens from the apex towards the base of an axis (Hacke et al. 2017), so a smaller or more distal section should carry narrower, more densely packed conduits, which is the winter pattern. Density and *D*ₕ, the traits with the largest winter effects, tracked stele area. Two observations argue against size as the whole explanation: lumen area fraction, the trait least related to size, did not change, and spring sections, slightly larger than fall ones, were no less dense. The winter shift is real in these sections but not yet evidence of a seasonal adjustment. Serial sections at measured distances from the union, with bud size recorded and both planes from the same bud, would settle it.

The spring decline in the xylem fraction of the stele (Figure 7E) is the more robust seasonal signal, because it occurred at both positions and arose outside the xylem ring, from growth of the rest of the stele. Living tissue, unlike the ring, can enlarge within weeks through rehydration and cambial activity: bud water content rose between winter and spring (Figure 1A), and vascular development resumes in the axis and the buds during deacclimation (Flinn & Ashworth 1994b; Savage & Chuine 2021; Kasuga et al. 2023). Its timing matches the maturation of the trace (Figure 6) and the loss of ice exclusion (Figure 4), so it marks the same transition at the scale of the whole axis.

Nor is the junction a constriction of the kind that isolates the reproductive shoots of *Calluna vulgaris*, whose pedicel-base ice barrier has up to 5.7 times less xylem area than vegetative shoots, all of it tracheids (Kuprian et al. 2016): the bud-base ring of *V. corymbosum* was at most about a third smaller than the stem ring and carried more conduits per unit area. If the junction contributes to ice exclusion, it does so through properties that counts and diameters do not capture, such as the pit-membrane pores implicated in the *Calluna* barrier, wall chemistry or hydration.

### Extraorgan freezing, water, and what remains open

The thermal and cryomicroscopic data agree with Flinn and Ashworth (1994a, b) on the strategy and extend their record in three ways. First, no LTE was detected to -46 °C in 42 buds across three seasons, including the hardy winter state. Second, the sequence of freezing is now timed: it began in the stem, consistent with the ice nucleators of the stem bark (Kishimoto et al. 2014) and as in cranberry (Villouta et al. 2022), reached the bud within minutes, and the bud froze slowly behind it, the thermal signature of extraorgan freezing (Ishikawa & Sakai 1985; Neuner et al. 2019a). Third, ice was imaged directly in intact frozen buds at -10 °C rather than inferred from voids left in tissue fixed while frozen, as in peach and *Forsythia* (Ashworth et al. 1989; Ashworth 1990) and in Flinn and Ashworth’s buds at -5 °C. The two records agree on the scales and the primordia but differ on the axis, where the earlier study found no voids in hardy buds and suggested that too little water had frozen to leave them. In our sections the bud axis looked frozen in every season without a discrete mass, which supports that explanation and places the barrier above the axis, between it and each flower primordium. Our late-October collection (LT_50_ -20 °C, primordia ice-free) matches their early acclimated state, not their nonacclimated September buds, killed at -5 °C with ice in the ovaries. In spring the primordia froze without a discrete ice mass, and in IDTA the whole bud froze almost at once with a larger exotherm, as a hydrated primordium reached by ice through a mature trace should. The fall-to-winter gain in hardiness occurred without a change in mean water content, so its most likely basis is cellular acclimation of the primordia, not measured here: in blueberry, dehydrins accumulate in floral buds during acclimation (Muthalif & Rowland 1994; Arora et al. 1997), a CBF transcription factor is expressed in floral buds in fall (Polashock et al. 2010), and soluble sugars accumulate in shoots (Lee et al. 2012). What changed over that interval was the distribution: the bimodal fall sample, with a dry subgroup that had no counterpart in winter, became uniform. Bulk dehydration thus appears to be an early, asynchronous component of acclimation, complete before the primordia reach their winter state, rather than the variable that sets midwinter hardiness.

The design is correlative, from one site and one annual cycle. The anatomical data rest on three buds per season, so they rely on exact tests, effect sizes whose intervals exclude zero, and a consistent direction across landmarks and traits. Only 27 % of elements had a readable wall pattern, fewest in the dormant seasons, and one observer scored one section per bud. Cryomicroscopy used a single temperature, and IDTA two seasons.

Three questions remain open. First, what kills the primordia while they exclude ice: failure of the barrier, or dehydration beyond what they tolerate (Kasuga et al. 2020; Villouta et al. 2020; Sakahara et al. 2025)? Second, does the trace mature before ice returns to the primordia, or are both outputs of one deacclimation program driven by chilling fulfillment and warmth (Kalberer et al. 2006; Kovaleski 2022; North et al. 2022)? Third, does hydration of the trace, rather than its wall pattern, let ice through? If maturation is part of the barrier, primordia should not freeze while the trace is immature, buds held in a hardy state should keep the immature pattern, and IDTA (Hacker & Neuner 2007) or cryomicroscopy at successive temperatures (Stegner et al. 2020) should show ice stopping below the primordia down to near the seasonal LT_50_. Wall-pattern scoring, which needs only a stained section, could then become a cheap marker of the loss of hardiness, useful in a crop whose spring primordia were injured from -8.5 °C (LT₁₀) and whose flower buds are damaged by late spring frosts (Hancock et al. 1987), a risk that warming may raise rather than lower (Gu et al. 2008; Kirchhof et al. 2025).

## Conclusions

Floral buds of *V. corymbosum* freeze extraorgan, and the barrier that keeps ice out of their flowers is not at the bud base: ice crossed the bud-stem junction and froze the bud axis in every season but stopped between the axis and each flower primordium. Work on ice barriers in this species should therefore move from the bud base, where the barrier lies in cranberry, to the pedicel and base of each primordium. The xylem trace along this path is present and lignified all year but mostly immature while the primordia exclude ice, and matures as exclusion is lost in spring. Its maturation, not its presence, tracks the loss of ice exclusion and, possibly with rehydration of the trace, is a candidate component of the barrier unexplained since 1994. The differentiation state of pre-formed vascular traces deserves attention wherever the spring reconnection of bud and stem is taken to explain frost exposure, and in this crop it could offer an anatomical marker of hardiness loss.

## Declarations

## Acknowledgements

The authors thank Faye Rosin, William Friedman and the whole team of the Arnold Arboretum of Harvard University for their support and for access to plant material.

## Author contributions

C.V.: conceptualization, methodology, investigation, formal analysis, data curation, visualization, writing (original draft), writing (review and editing), supervision, project administration and funding acquisition. A.K.R. and A.A.: investigation (sample processing and data collection).

## Data availability statement

The data that support the findings of this study are available from the corresponding author upon reasonable request.

## References

1. Andreini L., Viti R., Bartolini S., Ruiz D., Egea J. & Campoy J.A. (2012) The relationship between xylem differentiation and dormancy evolution in apricot flower buds (*Prunus armeniaca* L.): the influence of environmental conditions in two Mediterranean areas. Trees 26, 919–928. 10.1007/s00468-011-0668-1

2. Arora R. & Rowland L.J. (2011) Physiological research on winter-hardiness: deacclimation resistance, reacclimation ability, photoprotection strategies, and a cold acclimation protocol design. HortScience 46, 1070–1078. 10.21273/HORTSCI.46.8.1070

3. Arora R. & Taulavuori K. (2016) Increased risk of freeze damage in woody perennials vis-à-vis climate change: importance of deacclimation and dormancy response. Frontiers in Environmental Science 4, 44. 10.3389/fenvs.2016.00044

4. Arora R., Rowland L.J. & Panta G.R. (1997) Chill-responsive dehydrins in blueberry: are they associated with cold hardiness or dormancy transitions? Physiologia Plantarum 101, 8–16. 10.1111/j.1399-3054.1997.tb01813.x

5. Arora R., Rowland L.J., Ogden E.L., Dhanaraj A.L., Marian C.O., Ehlenfeldt M.K. & Vinyard B. (2004) Dehardening kinetics, bud development, and dehydrin metabolism in blueberry cultivars during deacclimation at constant, warm temperatures. Journal of the American Society for Horticultural Science 129, 667–674. 10.21273/JASHS.129.5.0667

6. Ashworth E.N. (1984) Xylem development in *Prunus* flower buds and the relationship to deep supercooling. Plant Physiology 74, 862–865. 10.1104/pp.74.4.862

7. Ashworth E.N. (1990) The formation and distribution of ice within *Forsythia* flower buds. Plant Physiology 92, 718–725. 10.1104/pp.92.3.718

8. Ashworth E.N. & Abeles F.B. (1984) Freezing behavior of water in small pores and the possible role in the freezing of plant tissues. Plant Physiology 76, 201–204. 10.1104/pp.76.1.201

9. Ashworth E.N., Davis G.A. & Wisniewski M.E. (1989) The formation and distribution of ice within dormant and deacclimated peach flower buds. Plant, Cell & Environment 12, 521–528. 10.1111/j.1365-3040.1989.tb02125.x

10. Ashworth E.N., Willard T.J. & Malone S.R. (1992) The relationship between vascular differentiation and the distribution of ice within *Forsythia* flower buds. Plant, Cell & Environment 15, 607–612. 10.1111/j.1365-3040.1992.tb01495.x

11. Bertel C., Hacker J. & Neuner G. (2021) Protective role of ice barriers: how reproductive organs of early flowering and mountain plants escape frost injuries. Plants 10, 1031. 10.3390/plants10051031

12. Biermann J., Stushnoff C. & Burke M.J. (1979) Differential thermal analysis and freezing injury in cold hardy blueberry flower buds. Journal of the American Society for Horticultural Science 104, 444–449. 10.21273/JASHS.104.4.444

13. Bittenbender H.C. & Howell G.S. (1974) Adaptation of the Spearman-Kärber method for estimating the T_50_ of cold stressed flower buds. Journal of the American Society for Horticultural Science 99, 187–190. 10.21273/JASHS.99.2.187

14. Bittenbender H.C. & Howell G.S. (1975) Interactions of temperature and moisture content on spring de-acclimation of flower buds of highbush blueberry. Canadian Journal of Plant Science 55, 447–452. 10.4141/cjps75-070

15. Bittenbender H.C. & Howell G.S. (1976) Cold hardiness of flower buds from selected highbush blueberry cultivars (*Vaccinium australe* Small). Journal of the American Society for Horticultural Science 101, 135–139. 10.21273/JASHS.101.2.135

16. Bond J., Donaldson L., Hill S. & Hitchcock K. (2008) Safranine fluorescent staining of wood cell walls. Biotechnic & Histochemistry 83, 161–171. 10.1080/10520290802373354

17. Burke M.J., Gusta L.V., Quamme H.A., Weiser C.J. & Li P.H. (1976) Freezing and injury in plants. Annual Review of Plant Physiology 27, 507–528. 10.1146/annurev.pp.27.060176.002451

18. Charrier G., Kovaleski A.P., Wenden B. & Hänninen H. (2026) Cold hardiness mechanisms and modeling: existing approaches and future avenues. New Phytologist 249, 2668–2682. 10.1111/nph.70863

19. Cromey D.W. (2010) Avoiding twisted pixels: ethical guidelines for the appropriate use and manipulation of scientific digital images. Science and Engineering Ethics 16, 639–667. 10.1007/s11948-010-9201-y

20. Cumming G. & Finch S. (2001) A primer on the understanding, use, and calculation of confidence intervals that are based on central and noncentral distributions. Educational and Psychological Measurement 61, 532–574. 10.1177/0013164401614002

21. Davis S.D., Sperry J.S. & Hacke U.G. (1999) The relationship between xylem conduit diameter and cavitation caused by freezing. American Journal of Botany 86, 1367–1372. 10.2307/2656919

22. Ehlenfeldt M.K., Rowland L.J., Ogden E.L. & Vinyard B.T. (2007) Floral bud cold hardiness of *Vaccinium ashei*, *V. constablaei*, and hybrid derivatives and the potential for producing northern-adapted rabbiteye cultivars. HortScience 42, 1131–1134. 10.21273/HORTSCI.42.5.1131

23. Eilers P.H.C. (2003) A perfect smoother. Analytical Chemistry 75, 3631–3636. 10.1021/ac034173t

24. Elzhov T.V., Mullen K.M., Spiess A.-N. & Bolker B. (2023) minpack.lm: R interface to the Levenberg-Marquardt nonlinear least-squares algorithm found in MINPACK, plus support for bounds. R package version 1.2–4. https://CRAN.R-project.org/package=minpack.lm

25. Endoh K., Kuwabara C., Arakawa K. & Fujikawa S. (2014) Consideration of the reasons why dormant buds of trees have evolved extraorgan freezing as an adaptation for winter survival. Environmental and Experimental Botany 106, 52–59. 10.1016/j.envexpbot.2014.02.008

26. Evert R.F. (2006) Esau’s Plant Anatomy: Meristems, Cells, and Tissues of the Plant Body: Their Structure, Function, and Development, 3rd edn. John Wiley & Sons, Hoboken, NJ. 10.1002/0470047380

27. Flinn C.L. & Ashworth E.N. (1994a) Blueberry flower-bud hardiness is not estimated by differential thermal analysis. Journal of the American Society for Horticultural Science 119, 295–298. 10.21273/JASHS.119.2.295

28. Flinn C.L. & Ashworth E.N. (1994b) Seasonal changes in ice distribution and xylem development in blueberry flower buds. Journal of the American Society for Horticultural Science 119, 1176–1184. 10.21273/JASHS.119.6.1176

29. George M.F., Burke M.J. & Weiser C.J. (1974) Supercooling in overwintering azalea flower buds. Plant Physiology 54, 29–35. 10.1104/pp.54.1.29

30. Gu L., Hanson P.J., Post W.M., Kaiser D.P., Yang B., Nemani R., … Meyers T. (2008) The 2007 eastern US spring freeze: increased cold damage in a warming world? BioScience 58, 253–262. 10.1641/B580311

31. Hacke U.G., Sperry J.S., Pockman W.T., Davis S.D. & McCulloh K.A. (2001) Trends in wood density and structure are linked to prevention of xylem implosion by negative pressure. Oecologia 126, 457–461. 10.1007/s004420100628

32. Hacke U.G., Spicer R., Schreiber S.G. & Plavcová L. (2017) An ecophysiological and developmental perspective on variation in vessel diameter. Plant, Cell & Environment 40, 831–845. 10.1111/pce.12777

33. Hacker J. & Neuner G. (2007) Ice propagation in plants visualized at the tissue level by infrared differential thermal analysis (IDTA). Tree Physiology 27, 1661–1670. 10.1093/treephys/27.12.1661

34. Hancock J.F., Nelson J.W., Bittenbender H.C., Callow P.W., Cameron J.S., Krebs S.L., … Schumann C.M. (1987) Variation among highbush blueberry cultivars in susceptibility to spring frost. Journal of the American Society for Horticultural Science 112, 702–706. 10.21273/JASHS.112.4.702

35. Hanson E.J., Berkheimer S.F. & Hancock J.F. (2007) Seasonal changes in the cold hardiness of the flower buds of highbush blueberry with varying species ancestry. Journal of the American Pomological Society 61, 14–18. 10.71318/apom.2007.61.1.14

36. Harris C.R., Millman K.J., van der Walt S.J., Gommers R., Virtanen P., Cournapeau D., … Oliphant T.E. (2020) Array programming with NumPy. Nature 585, 357–362. 10.1038/s41586-020-2649-2

37. Hedges L.V. (1981) Distribution theory for Glass’s estimator of effect size and related estimators. Journal of Educational Statistics 6, 107–128. 10.3102/10769986006002107

38. Houghton E., Watanabe Y., Neilsen D., Nelson L.M. & Hannam K. (2024) Investigating properties of sweet cherry (*Prunus avium*) flower buds that help promote freezing avoidance by supercooling. Plant Biology 26, 1067–1078. 10.1111/plb.13697

39. Ishikawa M. & Sakai A. (1981) Freezing avoidance mechanisms by supercooling in some *Rhododendron* flower buds with reference to water relations. Plant and Cell Physiology 22, 953–967. 10.1093/oxfordjournals.pcp.a076259

40. Ishikawa M. & Sakai A. (1985) Extraorgan freezing in wintering flower buds of *Cornus officinalis* Sieb. et Zucc. Plant, Cell & Environment 8, 333–338. 10.1111/j.1365-3040.1985.tb01407.x

41. Ishikawa M., Price W.S., Ide H. & Arata Y. (1997) Visualization of freezing behaviors in leaf and flower buds of full-moon maple by nuclear magnetic resonance microscopy. Plant Physiology 115, 1515–1524. 10.1104/pp.115.4.1515

42. Ishikawa M., Ide H., Yamazaki H., Murakawa H., Kuchitsu K., Price W.S. & Arata Y. (2016) Freezing behaviours in wintering *Cornus florida* flower bud tissues revisited using MRI. Plant, Cell & Environment 39, 2663–2675. 10.1111/pce.12813

43. Jones K.S., McKersie B.D. & Paroschy J. (2000) Prevention of ice propagation by permeability barriers in bud axes of *Vitis vinifera*. Canadian Journal of Botany 78, 3–9. 10.1139/b99-137

44. Kader S.A. & Proebsting E.L. (1992) Freezing behavior of *Prunus*, subgenus *Padus*, flower buds. Journal of the American Society for Horticultural Science 117, 955–960. 10.21273/JASHS.117.6.955

45. Kalberer S.R., Wisniewski M. & Arora R. (2006) Deacclimation and reacclimation of cold-hardy plants: current understanding and emerging concepts. Plant Science 171, 3–16. 10.1016/j.plantsci.2006.02.013

46. Kasuga J., Tsumura Y., Kondoh D., Jitsuyama Y., Horiuchi R. & Arakawa K. (2020) Cryo-scanning electron microscopy reveals that supercooling of overwintering buds of freezing-resistant interspecific hybrid grape ‘Yamasachi’ is accompanied by partial dehydration. Journal of Plant Physiology 253, 153248. 10.1016/j.jplph.2020.153248

47. Kasuga J., Kawase A., Kamigaki R., Mihara Y. & Kondoh D. (2023) Vessel development in compound buds of interspecific hybrid grape induced by artificial deacclimation treatments. Physiologia Plantarum 175, e13940. 10.1111/ppl.13940

48. Kirchhof E., Campos-Arguedas F., Arias N.S. & Kovaleski A.P. (2025) Thresholds for spring freeze: measuring risk to improve predictions in a warming world. New Phytologist 248, 563–575. 10.1111/nph.70453

49. Kishimoto T., Yamazaki H., Saruwatari A., Murakawa H., Sekozawa Y., Kuchitsu K., … Ishikawa M. (2014) High ice nucleation activity located in blueberry stem bark is linked to primary freeze initiation and adaptive freezing behaviour of the bark. AoB PLANTS 6, plu044. 10.1093/aobpla/plu044

50. Kogan C., DeVetter L.W. & Hoheisel G.-A. (2023) Modeling northern highbush blueberry cold hardiness for the Pacific Northwest. HortScience 58, 1314–1320. 10.21273/HORTSCI17320-23

51. Kovaleski A.P. (2022) Woody species do not differ in dormancy progression: differences in time to budbreak due to forcing and cold hardiness. Proceedings of the National Academy of Sciences of the United States of America 119, e2112250119. 10.1073/pnas.2112250119

52. Kovaleski A.P., Londo J.P. & Finkelstein K.D. (2019) X-ray phase contrast imaging of *Vitis* spp. buds shows freezing pattern and correlation between volume and cold hardiness. Scientific Reports 9, 14949. 10.1038/s41598-019-51415-2

53. Kuprian E., Briceño V.F., Wagner J. & Neuner G. (2014) Ice barriers promote supercooling and prevent frost injury in reproductive buds, flowers and fruits of alpine dwarf shrubs throughout the summer. Environmental and Experimental Botany 106, 4–12. 10.1016/j.envexpbot.2014.01.011

54. Kuprian E., Tuong T.D., Pfaller K., Wagner J., Livingston D.P. III & Neuner G. (2016) Persistent supercooling of reproductive shoots is enabled by structural ice barriers being active despite an intact xylem connection. PLoS ONE 11, e0163160. 10.1371/journal.pone.0163160

55. Kuprian E., Munkler C., Resnyak A., Zimmermann S., Tuong T.D., Gierlinger N., … Neuner G. (2017) Complex bud architecture and cell-specific chemical patterns enable supercooling of *Picea abies* bud primordia. *Plant*, Cell & Environment 40, 3101–3112. 10.1111/pce.13078

56. Lee J.H., Yu D.J., Kim S.J., Choi D. & Lee H.J. (2012) Intraspecies differences in cold hardiness, carbohydrate content and β-amylase gene expression of *Vaccinium corymbosum* during cold acclimation and deacclimation. Tree Physiology 32, 1533–1540. 10.1093/treephys/tps102

57. Lim C.C., Arora R. & Townsend E.C. (1998) Comparing Gompertz and Richards functions to estimate freezing injury in *Rhododendron* using electrolyte leakage. Journal of the American Society for Horticultural Science 123, 246–252. 10.21273/JASHS.123.2.246

58. Mayr S. & Sperry J.S. (2010) Freeze-thaw-induced embolism in *Pinus contorta*: centrifuge experiments validate the ‘thaw-expansion hypothesis’ but conflict with ultrasonic emission data. New Phytologist 185, 1016–1024. 10.1111/j.1469-8137.2009.03133.x

59. McMann N., Peichel A. & Savage J.A. (2022) Early spring flowers rely on xylem hydration but are not limited by stem xylem conductivity. New Phytologist 233, 838–850. 10.1111/nph.17782

60. Mills L.J., Ferguson J.C. & Keller M. (2006) Cold-hardiness evaluation of grapevine buds and cane tissues. American Journal of Enology and Viticulture 57, 194–200. 10.5344/ajev.2006.57.2.194

61. Muthalif M.M. & Rowland L.J. (1994) Identification of dehydrin-like proteins responsive to chilling in floral buds of blueberry (*Vaccinium*, section *Cyanococcus*). Plant Physiology 104, 1439–1447. 10.1104/pp.104.4.1439

62. Neuner G., Xu B. & Hacker J. (2010) Velocity and pattern of ice propagation and deep supercooling in woody stems of *Castanea sativa*, *Morus nigra* and *Quercus robur* measured by IDTA. Tree Physiology 30, 1037–1045. 10.1093/treephys/tpq059

63. Neuner G., Monitzer K., Kaplenig D. & Ingruber J. (2019a) Frost survival mechanism of vegetative buds in temperate trees: deep supercooling and extraorgan freezing vs. ice tolerance. Frontiers in Plant Science 10, 537. 10.3389/fpls.2019.00537

64. Neuner G., Kreische B., Kaplenig D., Monitzer K. & Miller R. (2019b) Deep supercooling enabled by surface impregnation with lipophilic substances explains the survival of overwintering buds at extreme freezing. Plant, Cell & Environment 42, 2065–2074. 10.1111/pce.13545

65. North M.G., Workmaster B.A. & Atucha A. (2022) Effects of chill unit accumulation and temperature on woody plant deacclimation kinetics. Physiologia Plantarum 174, e13717. 10.1111/ppl.13717

66. Pearce R.S. & Fuller M.P. (2001) Freezing of barley studied by infrared video thermography. Plant Physiology 125, 227–240. 10.1104/pp.125.1.227

67. Pittermann J. & Sperry J.S. (2003) Tracheid diameter is the key trait determining the extent of freezing-induced embolism in conifers. Tree Physiology 23, 907–914. 10.1093/treephys/23.13.907

68. Pittermann J. & Sperry J.S. (2006) Analysis of freeze-thaw embolism in conifers. The interaction between cavitation pressure and tracheid size. Plant Physiology 140, 374–382. 10.1104/pp.105.067900

69. Polashock J.J., Arora R., Peng Y., Naik D. & Rowland L.J. (2010) Functional identification of a C-repeat binding factor transcriptional activator from blueberry associated with cold acclimation and freezing tolerance. Journal of the American Society for Horticultural Science 135, 40–48. 10.21273/JASHS.135.1.40

70. Quamme H.A., Su W.A. & Veto L.J. (1995) Anatomical features facilitating supercooling of the flower within the dormant peach flower bud. Journal of the American Society for Horticultural Science 120, 814–822. 10.21273/JASHS.120.5.814

71. R Core Team (2023) R: A Language and Environment for Statistical Computing. R Foundation for Statistical Computing, Vienna, Austria. https://www.R-project.org/

72. Rajashekar C.B. (1989) Supercooling characteristics of isolated peach flower bud primordia. Plant Physiology 89, 1031–1034. 10.1104/pp.89.4.1031

73. Rodrigo J. (2000) Spring frosts in deciduous fruit trees: morphological damage and flower hardiness. Scientia Horticulturae 85, 155–173. 10.1016/S0304-4238(99)00150-8

74. Rowland L.J., Ogden E.L., Ehlenfeldt M.K. & Vinyard B. (2005) Cold hardiness, deacclimation kinetics, and bud development among 12 diverse blueberry genotypes under field conditions. Journal of the American Society for Horticultural Science 130, 508–514. 10.21273/JASHS.130.4.508

75. Rowland L.J., Ogden E.L., Ehlenfeldt M.K. & Arora R. (2008) Cold tolerance of blueberry genotypes throughout the dormant period from acclimation to deacclimation. HortScience 43, 1970–1974. 10.21273/HORTSCI.43.7.1970

76. Sakahara S., Atucha A. & Kasuga J. (2025) Freeze-induced dehydration and its impact on freezing resistance in dormant grapevine buds. Physiologia Plantarum 177, e70682. 10.1111/ppl.70682

77. Sakai A. & Larcher W. (1987) Frost Survival of Plants: Responses and Adaptation to Freezing Stress. Ecological Studies 62. Springer-Verlag, Berlin. 10.1007/978-3-642-71745-1

78. Savage J.A. & Chuine I. (2021) Coordination of spring vascular and organ phenology in deciduous angiosperms growing in seasonally cold climates. New Phytologist 230, 1700–1715. 10.1111/nph.17289

79. Schneider C.A., Rasband W.S. & Eliceiri K.W. (2012) NIH Image to ImageJ: 25 years of image analysis. Nature Methods 9, 671–675. 10.1038/nmeth.2089

80. Scholz A., Klepsch M., Karimi Z. & Jansen S. (2013) How to quantify conduits in wood? Frontiers in Plant Science 4, 56. 10.3389/fpls.2013.00056

81. Signorelli S., Shaw J., Hermawaty D., Wang Z., Verboven P., Considine J.A. & Considine M.J. (2020) The initiation of bud burst in grapevine features dynamic regulation of the apoplastic pore size. Journal of Experimental Botany 71, 719–729. 10.1093/jxb/erz200

82. Sperry J.S. & Sullivan J.E.M. (1992) Xylem embolism in response to freeze-thaw cycles and water stress in ring-porous, diffuse-porous, and conifer species. Plant Physiology 100, 605–613. 10.1104/pp.100.2.605

83. Sperry J.S., Nichols K.L., Sullivan J.E.M. & Eastlack S.E. (1994) Xylem embolism in ring-porous, diffuse-porous, and coniferous trees of northern Utah and interior Alaska. Ecology 75, 1736–1752. 10.2307/1939633

84. Stegner M., Wagner J. & Neuner G. (2020) Ice accommodation in plant tissues pinpointed by cryo-microscopy in reflected-polarised-light. Plant Methods 16, 73. 10.1186/s13007-020-00617-1

85. Takahashi D., Willick I.R., Kasuga J. & Livingston D.P. III (2021) Responses of the plant cell wall to sub-zero temperatures: a brief update. Plant and Cell Physiology 62, 1858–1866. 10.1093/pcp/pcab103

86. Villouta C., Workmaster B.A., Bolivar-Medina J., Sinclair S. & Atucha A. (2020) Freezing stress survival mechanisms in *Vaccinium macrocarpon* Ait. terminal buds. Tree Physiology 40, 841–855. 10.1093/treephys/tpaa028

87. Villouta C., Workmaster B.A. & Atucha A. (2021) Freezing stress damage and growth viability in *Vaccinium macrocarpon* Ait. bud structures. Physiologia Plantarum 172, 2238–2250. 10.1111/ppl.13457

88. Villouta C., Workmaster B.A., Livingston D.P. III & Atucha A. (2022) Acquisition of freezing tolerance in *Vaccinium macrocarpon* Ait. is a multi-factor process involving the presence of an ice barrier at the bud base. Frontiers in Plant Science 13, 891488. 10.3389/fpls.2022.891488

89. Virtanen P., Gommers R., Oliphant T.E., Haberland M., Reddy T., Cournapeau D., … SciPy 1.0 Contributors (2020) SciPy 1.0: fundamental algorithms for scientific computing in Python. Nature Methods 17, 261–272. 10.1038/s41592-019-0686-2

90. Wickham H. (2016) ggplot2: Elegant Graphics for Data Analysis, 2nd edn. Springer, Cham. 10.1007/978-3-319-24277-4

91. Wisniewski M., Davis G. & Schafter K. (1991) Mediation of deep supercooling of peach and dogwood by enzymatic modifications in cell-wall structure. Planta 184, 254–260. 10.1007/BF00197955

92. Wisniewski M., Lindow S.E. & Ashworth E.N. (1997) Observations of ice nucleation and propagation in plants using infrared video thermography. Plant Physiology 113, 327–334. 10.1104/pp.113.2.327

93. Wisniewski M., Gusta L. & Neuner G. (2014) Adaptive mechanisms of freeze avoidance in plants: a brief update. Environmental and Experimental Botany 99, 133–140. 10.1016/j.envexpbot.2013.11.011

94. Wisniewski M., Neuner G. & Gusta L.V. (2015) The use of high-resolution infrared thermography (HRIT) for the study of ice nucleation and ice propagation in plants. Journal of Visualized Experiments 99, e52703. 10.3791/52703

95. Workmaster B.A.A., Palta J.P. & Wisniewski M. (1999) Ice nucleation and propagation in cranberry uprights and fruit using infrared video thermography. Journal of the American Society for Horticultural Science 124, 619–625. 10.21273/JASHS.124.6.619

96. Zohner C.M., Mo L., Renner S.S., Svenning J.-C., Vitasse Y., Benito B.M., … Crowther T.W. (2020) Late-spring frost risk between 1959 and 2017 decreased in North America but increased in Europe and Asia. Proceedings of the National Academy of Sciences 117, 12192–12200. 10.1073/pnas.1920816117

